# Specialized and shared functions of diguanylate cyclases and phosphodiesterases in *Streptomyces* development

**DOI:** 10.1101/2020.04.24.060822

**Authors:** Julian Haist, Sara Alina Neumann, Mahmoud M Al-Bassam, Sandra Lindenberg, Marie A Elliot, Natalia Tschowri

**Author notes:** GlaxoSmithKline GmbH & Co. KG, Prinzregentenplatz 9, 81675 München, Germany. These authors contributed equally to this work (Julian Haist, Sara Alina Neumann).

## Abstract

Levels of the second messenger bis-3’-5’-cyclic di-guanosinemonophosphate (c-di-GMP) determine when *Streptomyces* initiate sporulation to survive under adverse conditions. c-di-GMP signals are integrated into the genetic differentiation network by the regulator BldD and the sigma factor σ^WhiG^. However, functions of the development-specific c-di-GMP diguanylate cyclases (DGCs) CdgB and CdgC, and the phosphodiesterases (PDEs) RmdA and RmdB, are poorly understood. Here, we provide biochemical evidence that the GGDEF-EAL domain protein RmdB from *S. venezuelae* is a monofunctional PDE that hydrolyzes c-di-GMP to 5’pGpG. Despite having an equivalent GGDEF-EAL domain arrangement, RmdA cleaves c-di-GMP to GMP and exhibits residual DGC activity. We show that an intact EAL motif is crucial for the *in vivo* function of both enzymes since strains expressing protein variants with an AAA motif instead of EAL are delayed in development, similar to null mutants. Global transcriptome analysis of Δ*cdgB*, Δ*cdgC*, Δ*rmdA* and Δ*rmdB* strains revealed that the c-di-GMP specified by these enzymes has a global regulatory role, with about 20 % of all *S. venezuelae* genes being differentially expressed in the *cdgC* mutant. Our data suggest that the major c-di-GMP-controlled targets determining the timing and mode of sporulation are genes involved cell division and the production of the hydrophobic sheath that covers *Streptomyces* aerial hyphae and spores. Altogether, this study provides a global view of the c-di-GMP-dependent genes that contribute to the hyphae-to-spores transition and sheds light on the shared and specific functions of the key enzymes involved in c-di-GMP metabolism in *S. venezuelae*.

**Importance:** *Streptomyces* are important producers of clinical antibiotics. The ability to synthesize these natural products is connected to their developmental biology, which includes a transition from filamentous cells to spores. The widespread bacterial second messenger c-di-GMP controls this complex switch and is a promising tool to improve antibiotic production. Here, we analyzed the enzymes that make and break c-di-GMP in *S. venezuelae* by studying the genome-wide transcriptional effects of the DGCs CdgB and CdgC and the PDEs RmdA and RmdB. We found that the c-di-GMP specified by these enzymes has a global regulatory role. However, despite shared enzymatic activities, the four c-di-GMP enzymes have specialized inputs into differentiation. Altogether, we demonstrate that altering c-di-GMP levels through the action of selected enzymes yields characteristically distinct transcriptional profiles; this can be an important consideration when modulating c-di-GMP for the purposes of natural product synthesis in *Streptomyces*.

## Introduction

Cellular levels of the bacterial second messenger bis-3’-5’-cyclic di-guanosinemonophosphate are controlled by the competing activities of GGDEF-domain containing diguanylate cyclases (DGCs) that produce the molecule out of GTP, and by phosphodiesterases (PDEs) that carry an EAL or HD-GYP domain to degrade the second messenger (1). Multiplicity of c-di-GMP-turnover genes within a genome is widespread in bacteria, making it challenging to understand how individual DGCs and PDEs control specific cellular responses while sharing common enzymatic activities (2). For example, *Escherichia coli* K-12 has 12 DGCs and 13 PDEs. While deleting distinct DGCs and PDEs has no effect on cellular c-di-GMP levels, it has drastic consequences for *E. coli* biofilm formation (3). In *Vibrio cholerae*, which possesses 53 proteins with c-di-GMP-metabolizing domains, only a subset of these proteins affects motility, biofilm formation, or both (4).

c-di-GMP is renowned for its function in guiding the transition between motility and sessility in most bacteria (5). High levels of the second messenger favor the switch to sessility, a process that often involves formation of self-organized, structured biofilms as a stationary-phase induced survival strategy. In the non-motile streptomycetes, c-di-GMP is a key factor controlling the transition between their filamentous lifestyle and spore formation. However, in these bacteria, *low* levels of the molecule favor initiation of their sporulation survival strategy. For example, overexpression of the *E. coli* PDE PdeH in *S. venezuelae* induces premature, massive sporulation (6). A classical *Streptomyces* life cycle includes the erection of hyphae into the air when the bacteria switch to their stationary growth phase, followed by the morphogenesis of these aerial filaments into chains of spores. In *S. venezuelae*, aerial mycelium formation is completely bypassed when c-di-GMP levels are too low, due to PDE overexpression (6). A phenotypically identical response can be caused through deleting the DGC-encoding gene *cdgC* – one of the 10 chromosomally-encoded GGDEF/EAL/HD-GYP genes in *S. venezuelae* (7, 8). Deletion of yet another DGC-encoding gene, *cdgB*, also leads to precocious sporulation; however, the *cdgB* mutant still undergoes the classical *Streptomyces* life cycle and forms spores on reproductive aerial hyphae like the wild type. Therefore, deleting *cdgB* shifts sporulation timing but does not affect the principle mode of spore formation, i.e. transition of aerial hyphae into chains of spores. Conversely, overexpressing the *S. coelicolor* DGC, CdgB, causes an opposing phenotype in *S. venezuelae*, in that it prolongs filamentous, vegetative growth (8); this process can be mimicked by deleting either *rmdA* or *rmdB*, which encode functional PDEs (9) (Fig. 1).

**Figure 1:**
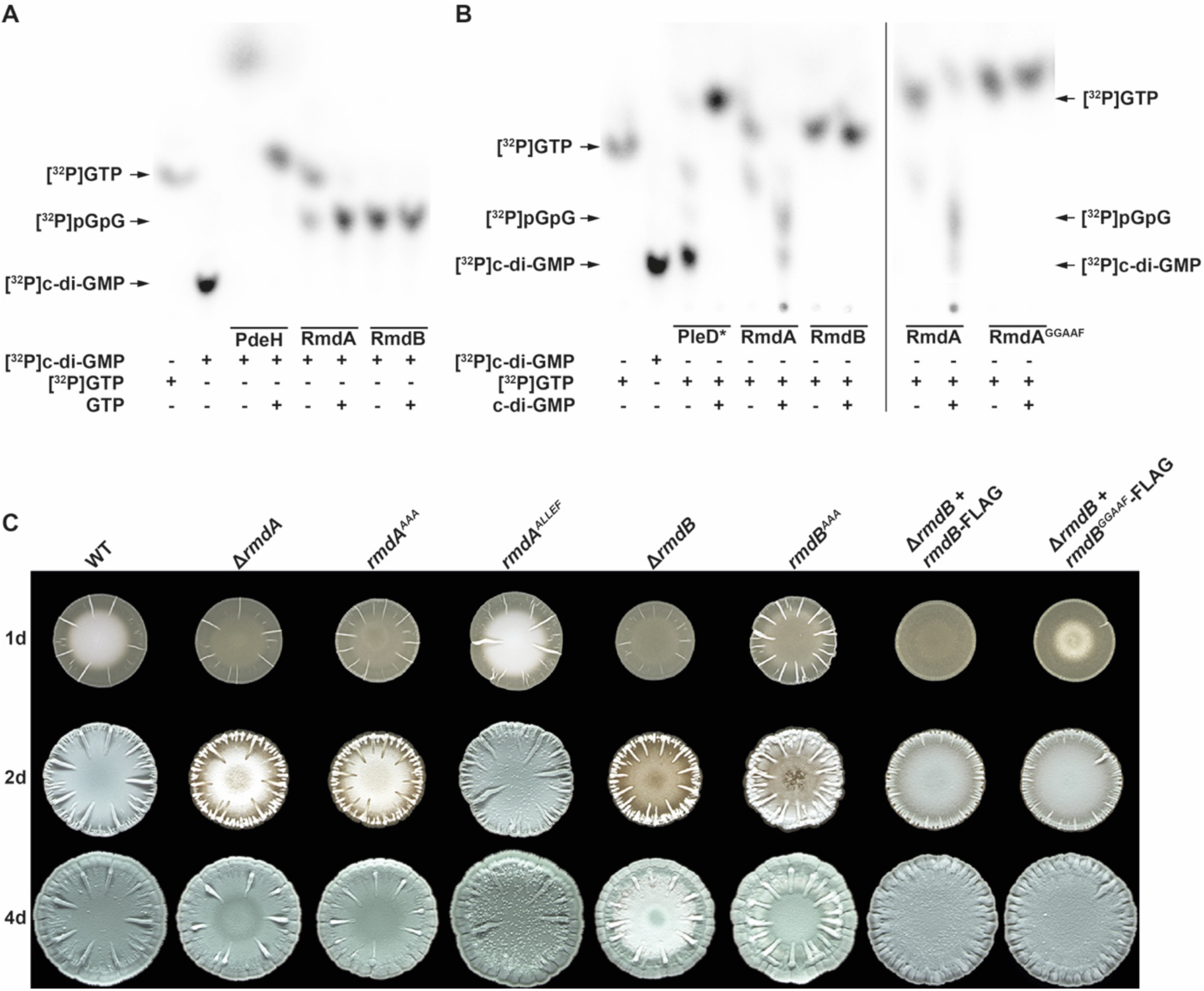
Enzymatic activities of RmdA and RmdB. Purified RmdA and RmdB were assayed for PDE activity with [^32^P]c-di-GMP (2.08 nM) (A) and for DGC activity with [^32^P]GTP (4.16 nM) (B) as substrate. RmdA carrying a mutagenized GGDEF motif (RmdA^GGAAF^) was tested for DGC function in (B). Where indicated, samples also contained either 1 mM GTP or 1 mM c-di-GMP as competitors. The PDE PdeH from *E. coli* and the DGC PleD* from *C. crescentus* served as positive controls for the PDE and DGC assays, respectively. One µM of each purified protein was used in reactions. (C) Macrocolonies of *S. venezuelae* wild type and mutants were grown for up to 4 days (d) at 30 °C. Development of strains carrying the mutagenized AAA motif instead of EAL in the chromosomal *rmdA* (*rmdA*^AAA^) or *rmdB* (*rmdB*^AAA^) and corresponding deletion mutants (Δ*rmdA* or Δ*rmdB*) was analyzed. Wild type RmdB-FLAG and a mutant variant with the GGAAF motif instead of GGDEF (RmdB^GGAAF^-FLAG) were expressed from the Φ_BT1_ phage integration site under the control of the native *rmdB* promoter.

*Streptomyces* development is controlled by a complex network of Bld and Whi regulators. Strains mutated in *bld* genes fail to develop aerial hyphae, while deletion of *whi* genes blocks the transition of aerial hyphae into spores (10). c-di-GMP signals are integrated in the two *Streptomyces* cell-fate establishing stages and determine (I) the period of vegetative, filamentous growth by binding to the transcriptional regulator, BldD, (6) and (II) initiation of sporulation by controlling the activity of the sporulation-specific sigma factor σ^WhiG^ (11). Binding of c-di-GMP to BldD induces protein dimerization and stimulates BldD-binding to target DNA. In *S. venezuelae*, BldD binds to 282 target sequences *in vivo* and represses sporulation. Consequently, the *bldD* mutant bypasses aerial mycelium formation and sporulates precociously (6). σ^WhiG^ activity is determined by the anti-σ factor RsiG, which sequesters σ^WhiG^ when in complex with c-di-GMP. Upon release at low c-di-GMP levels, σ^WhiG^ directly activates three genes: *whiI, whiH* and *vnz1500*5. Through the sporulation-specific regulators WhiI and WhiH, σ^WhiG^ thus controls a large regulon of sporulation genes. Overexpressing either *whiG* or co-overexpressing *whiI* and *whiH*, induces hypersporulation (11), as seen for Δ*cdgC*.

Although direct targets of the known c-di-GMP-sensors in *Streptomyces*, BldD and σ^WhiG^, are known, there is a major gap in our understanding of c-di-GMP-responsive genes in the genus. The specific molecular targets driving hypersporulation in the two DGC mutants (Δ*cdgB* and Δ*cdgC*) on the one hand, and delaying sporulation in the two PDE mutants (Δ*rmdA* and Δ*rmdB*) on the other hand, are not defined. Moreover, it is unclear why the *cdgB* mutant forms premature spores within aerial hyphae, while the *cdgC* mutant is unable to raise aerial mycelium, despite their products possessing the same enzymatic function. To understand the shared, and specialized roles of the two DGCs and PDEs, respectively, we used RNA-sequencing (RNA-seq) to compare the transcriptional profiles of Δ*cdgB*, Δ*cdgC*, Δ*rmdA* and Δ*rmdB*, with wild type *S. venezuelae*. We found that expression of the hydrophobic sheath genes is strongly responsive to DGC and PDE deletions. Chaplin and rodlins are upregulated in Δ*cdgB*, but are downregulated in Δ*cdgC*, explaining the failure of Δ*cdgC* to raise aerial mycelium. Moreover, we show that DGCs and PDEs antagonistically control expression of cell division components, likely contributing to c-di-GMP-induced shifts in timing of sporulation initiation. The distinct regulons of the DGCs CdgB and CdgC, and of the PDEs RmdA and RmdB, imply that these enzymes orchestrate distinct cellular responses to specific environmental and metabolic signals.

## Results and discussion

### Biochemical and physiological activities of the GGDEF and EAL domains of RmdA and RmdB

The cytosolic RmdA and the membrane-bound RmdB are composite GGDEF-EAL domain proteins that are functional PDEs in *S. coelicolor* (9) but their enzymatic activities have not been characterized for the *S. venezuelae* homologs. The GGDEF and the EAL domains are fully conserved in both proteins. GGDEF domains bind GTP and can allosterically modulate the activities of EAL-domains when organized in tandem, as demonstrated for the GGDEF-EAL PDE CC3396 from *Caulobacter crescentus* (12). We wondered whether the GGDEF domains of RmdA and RmdB were capable of GTP conversion into c-di-GMP, or if they had any influence on the activity of their associated EAL domains. We purified RmdA fused to a maltose-binding protein (MBP) tag at its N-terminus, and an N-terminally 6×His-tagged cytosolic fraction of RmdB. The PDE PdeH from *Escherichia coli* and the DGC PleD* from *C. crescentus* served as positive controls for the PDE and DGC assays, respectively (13, 14). [^32^P]-labeled c-di-GMP or [^32^P]GTP was added as a substrate for *in vitro* PDE and DGC assays, respectively, and the reactions were separated by thin layer chromatography (TLC).

Our data show that RmdB hydrolyzed [^32^P]c-di-GMP to the linear [^32^P]pGpG (Fig. 1A). This reaction was more efficient in presence of manganese than magnesium ions, revealing that Mn^2+^ is the preferred cofactor (Fig. S1). In contrast, RmdA cleaved [^32^P]c-di-GMP to [^32^P]GMP via the intermediate [^32^P]pGpG. Interestingly, excess GTP inhibited the RmdA-mediated hydrolysis of [^32^P]pGpG to [^32^P]GMP, suggesting that GTP binding to the GGDEF domain compromises the PDE activity of the EAL domain (Fig. 1A).

Incubation of RmdB with [^32^P]GTP did not result in any reaction products, suggesting that the GGDEF domain is inactive, at least under the conditions tested here (Fig. 1B). In contrast, we detected an additional spot after separating the reaction sample containing RmdA and [^32^P]GTP. We hypothesized that this spot represented an intermediate product of c-di-GMP synthesis. To reduce the immediate EAL-domain-mediated hydrolysis of any [^32^P]c-di-GMP produced by the GGDEF domain of RmdA, we added non-labeled c-di-GMP as competitor. Indeed, we detected both [^32^P]c-di-GMP synthesized by RmdA, and [^32^P]pGpG that arose due to rapid degradation of [^32^P]c-di-GMP by its EAL domain (Fig. 1B). To confirm that c-di-GMP production by RmdA required an intact GGDEF site, we mutagenized the GGDEF to GGAAF motif and used purified MBP-RmdA^GGAAF^ in the DGC assays. As expected, neither c-di-GMP nor pGpG were detectable in the reaction containing the mutagenized RmdA^GGAAF^ protein (Fig. 1B). Altogether, these data show that RmdB from *S. venezuelae* is a monofunctional PDE that cleaves c-di-GMP to the linear pGpG. Conversely, RmdA hydrolyzes c-di-GMP to GMP via pGpG and has weak DGC activity that likely remains cryptic, since the c-di-GMP generated by the GGDEF domain appears to be immediately hydrolyzed by the PDE activity of the EAL domain. Such residual DGC activity in tandem proteins is not uncommon and has also been reported for the GGDEF-EAL protein PdeR from *E. coli* (15). However, we cannot exclude the possibility that, under certain conditions, the DGC activity of RmdA becomes dominant over the PDE function.

To assess the impact of the individual GGDEF and EAL domains of RmdA and RmdB on developmental control *in vivo*, we generated strains carrying chromosomal mutations in either GGDEF or EAL active sites. The strain expressing *rmdA* with an AAA motif instead of the EAL motif (*rmdA*^AAA^) showed a delay in development, similar to that of the *rmdA* mutant strain (Fig. 1C). In contrast, mutagenizing the GGDEF motif to ALLEF in the chromosomal locus of *rmdA* (*rmdA*^ALLEF^) had no effect on differentiation compared to wild type (Fig. 1C). Similarly, a strain carrying the mutant AAA motif (in place of the EAL motif) in the EAL domain of *rmdB* (*rmdB*^AAA^) was delayed in development, like the *rmdB* null mutant. We were unable to generate a strain expressing the *rmdB*^ALLEF^ allele from the chromosome, so instead we applied complementation analysis. We found that an *rmdB* allele carrying the mutagenized GGAAF motif in the GGDEF site could complement the differentiation defect of the *rmdB* mutant (Fig. 1C). These data suggest that a functional EAL domain is crucial for the *in vivo* functions of RmdA and RmdB. While the GGDEF domain of RmdA can synthesize c-di-GMP *in vitro*, this activity does not seem to contribute to differentiation control by RmdA *in vivo* under the conditions tested.

### Genome-wide transcriptional profiling of *S. venezuelae* c-di-GMP mutants

Controlling developmental progression is the key function of c-di-GMP in all tested *Streptomyces* models (6, 16-18). Nevertheless, targets of the second messenger have not yet been addressed on a genome-wide scale. Out of the ten GGDEF/EAL/HD-GYP-proteins encoded by *S. venezuelae*, only four enzymes control c-di-GMP-mediated differentiation processes. Deleting the DGC-encoding *cdgB* and *cdgC* causes precocious sporulation, but, the phenotypes of the two mutants differ, in that the Δ*cdgB* strain forms spores on aerial hyphae, whereas the Δ*cdgC* strain completely skips the aerial mycelium formation stage. On the other hand, deleting either the PDE-encoding *rmdA* or *rmdB* delays development. However, the phenotypes of these two strains are not identical: losing *rmdB* arrests *S. venezuelae* in the vegetative growth phase for ca. 1 day longer than deleting *rmdA* (8). These phenotypes suggest that the two DGCs and PDEs share common functions but also play unique roles in developmental regulation. To understand their functions, we conducted RNA-sequencing (RNA-seq) of the transcriptomes of the four mutants and their wild type parent strain. The distinct phenotypes of the Δ*cdgB*, Δ*cdgC*, Δ*rmdA* and Δ*rmdB* mutants were particularly pronounced when *S. venezuelae* was grown on Maltose-Yeast Extract-Malt Extract (MYM) agar. Hence, for the RNA-seq analyses, we harvested macrocolonies from plates that were inoculated with identical numbers of spores (12 µl of 2×10^5^ CFU/µl) and were grown for 30 hours at 30 °C. For each strain, three independent macrocolonies were pooled for RNA-isolation and two samples were sequenced per strain. Thus, the resulting transcriptional profiles would be representative of six (combined) biological replicates. At the time of harvest, wild type, Δ*rmdA* and Δ*rmdB* were at a vegetative stage of growth, while Δ*cdgB* and Δ*cdgC* had already sporulated (Fig. S2).

We were specifically interested in genes that are known components of cascades controlling differentiation (10); however, a complete table of differentially expressed genes is presented in Dataset S1. Genes that exhibited a more than 2-fold (log2 >1/<-1; p<0.05) increase or decrease in expression in the mutants relative to the wild type were considered as significant. Impressively, in the *cdgC* mutant, 1458 genes exhibited significant changes in transcription, with 844 genes being up-and 616 downregulated in comparison to the wild type (Fig. 2A, Dataset 1). In Δ*cdgB*, Δ*rmdA* and Δ*rmdB*, 312, 293 and 164 genes, respectively, were differentially expressed (Fig. 2A). Thus, we uncovered a broad regulon of c-di-GMP in *S. venezuelae* with CdgC affecting ∼20 % of all *S. venezuelae* open reading frames. We conclude that c-di-GMP controlled by CdgB, CdgC, RmdA and RmdB has a global regulatory role. The situation is different, for example, in *E. coli*, where the DGC DgcM and the PDE PdeR, antagonistically controlling the biosynthesis of adhesive curli-fibers, act in a precise and non-global manner on few specific targets (19).

**Figure 2:**
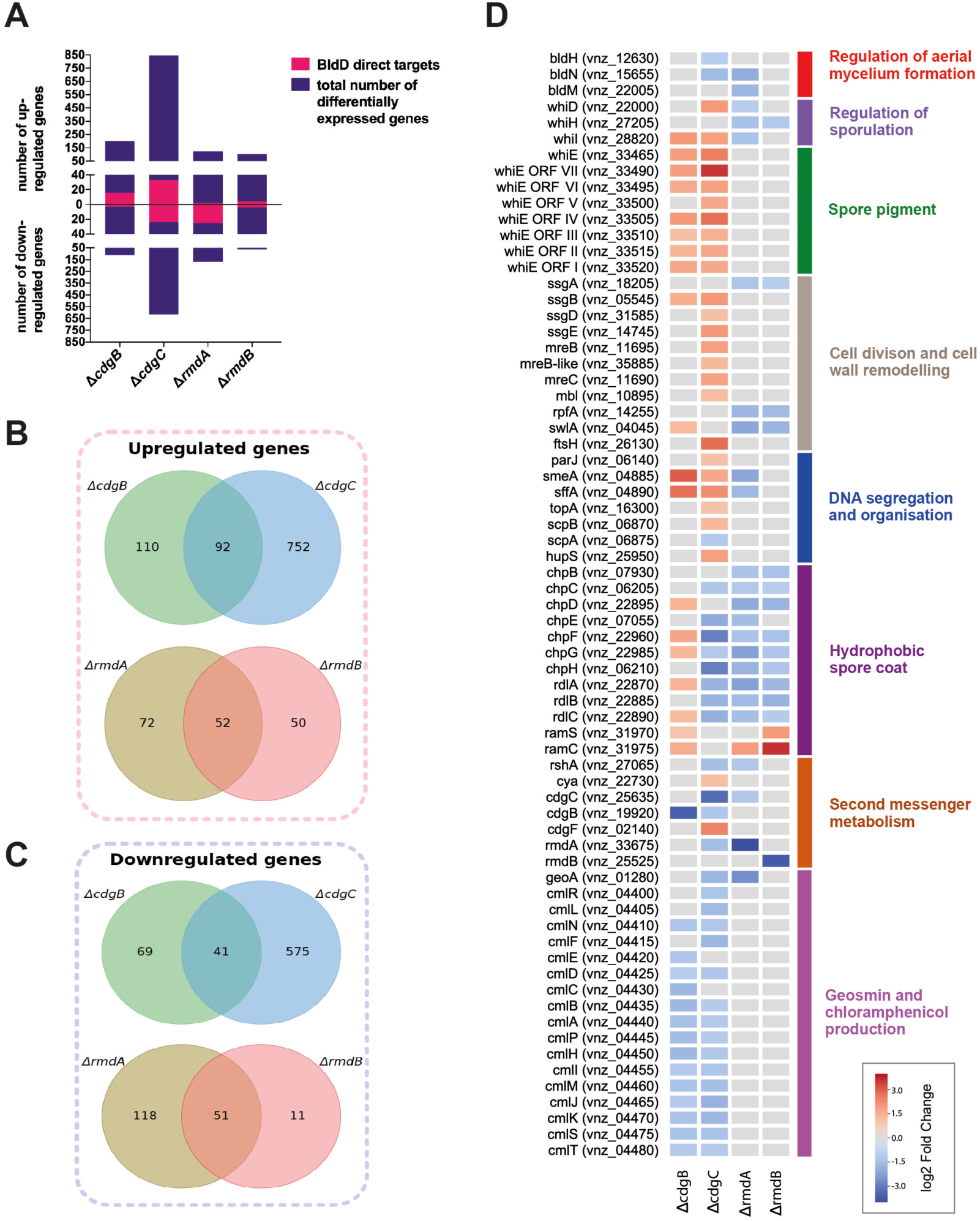
RNA-seq profiles of Δ*cdgB*, Δ*cdgC*, Δ*rmdA* and Δ*rmdB* strains. (A) Histogram showing total numbers of down- and upregulated genes in the analyzed mutants. Out of these, numbers of differentially expressed direct BldD-targets in each strain are visualized in pink. Venn diagrams depict number of upregulated (B) and downregulated (C) genes that overlap between the two DGC mutants Δ*cdgC* and Δ*cdgB*, or between the two PDE mutants Δ*rmdA* and Δ*rmdB*. (D) Heat map showing differentially expressed genes associated with developmental processes. Genes with 2-fold (log2 >1/<-1; p<0.05) change in expression were considered as significant.

By comparing the transcriptomes of Δ*cdgB* and Δ*cdgC* we found only 92 upregulated and 41 downregulated genes that were shared in the two DGC mutants (Fig. 2B and C). Thus, out of the 1770 genes that are in sum differentially expressed in the two mutants, only ∼8 % of genes overlapped. When examining the transcription profiles of the Δ*rmdA* and Δ*rmdB* mutants, we found 52 upregulated genes and 51 downregulated genes that were common to both strains (Fig. 2B and C). In total, this corresponds to ∼23 % of all differentially expressed genes being similarly impacted by both RmdA and RmdB. This shows that despite a shared enzymatic activity, the DGCs and the PDEs, respectively, control characteristic sets of genes. The N-termini of CdgC and RmdB are anchored in the cell membrane, CdgB has GAF-PAS-PAC N-terminal sensory domains and RmdA contains PAS-PAC domains at the N-terminus (7). Likely, the signals perceived by the characteristic sensory domains specify the distinct functions of CdgB, CdgC, RmdA and RmdB.

### *bld* and *whi* genes with altered expression in the DGC / PDE mutants

Proteins of the Bld and Whi families are key regulators of the developmental genetic network. BldD sits on top of the developmental regulatory cascade, and when in complex with c-di-GMP, it binds to 282 target promoters in the *S. venezuelae* chromosome (6, 10). BldD acts as a transcriptional repressor on most target promoters (20, 21), but it can also activate gene expression (18). Unexpectedly, we found few *bld* and *whi* genes to be differentially expressed in the studied mutants. In agreement with a delay in development, *bldN, bldM, whiD, whiH* and *whiI* were downregulated in Δ*rmdA*; however, of these, only *whiH* was also downregulated in Δ*rmdB*. In Δ*cdgB*, only *whiI* was upregulated at the tested time-point, while in Δ*cdgC*, both *whiI* and *whiD* were upregulated, while *bldH* and *bldN* were downregulated (Fig. 2D and S3A).

The expression of *whiI* and *whiH* is directly activated by the sigma factor σ^WhiG^, whose activity is controlled by the RsiG-(c-di-GMP) anti-sigma factor. Expression of *whiI* completely depends on *whiG*, whereas *whiH* expression is only partially dependent on the sigma factor (11). Thus, the fact that *whiI* was upregulated in both Δ*cdgB* and Δ*cdgC*, reflected the activation of σ^WhiG^ in the two DGC mutants. *whiH* and *whiI* were, however, both downregulated in Δ*rmdA; whiH* was also less expressed in Δ*rmdB.* This collectively suggests reduced activity of σ^WhiG^ in the two PDE mutant strains. The inversely correlated transcription profiles of these σ^WhiG^-dependent genes imply that the two DGCs and two PDEs all contribute to modulating σ^WhiG^-activity.

The BldN ECF sigma factor activates the expression of the chaplin and rodlin genes, which encode the hydrophobic sheath proteins that encase aerial hyphae and spores (22). BldD-(c-di-GMP) directly represses *bldN* expression (23) (18, 21). Thus, we expected increased transcription of *bldN* in the DGC mutants, due to loss of BldD repressive activities, and reduced expression of *bldN* in the PDE mutants. It was, therefore, a surprise that *bldN* expression was downregulated in the Δ*cdgC* strain. Because of that we set out to examine the expression patterns of all known BldD-(c-di-GMP) target genes in our different mutants. Of the 282 direct BldD-(c-di-GMP) targets in *S. venezuelae*, we found 19, 57, 27 and 8 genes to be differentially expressed in Δ*cdgB*, Δ*cdgC*, Δ*rmdA* and Δ*rmdB*, respectively (Fig. 2A).

These analyses revealed that, at least under the conditions tested, only a relatively minor fraction of all BldD-(c-di-GMP)-targets responded to c-di-GMP changes in the studied mutants. Notably, the direct BldD-regulon was determined in *S. venezuelae* grown in liquid culture, and some of the observed differences may be explained by the fact that here, colonies grown on solid medium were analyzed. However, many direct BldD-targets are co-regulated by multiple transcription factors in a hierarchical manner (10), and thus require multiple, additional signals for proper expression. For example, in *S. venezuelae*, the response regulator MtrA, binds directly to a number of *bld* and *whi* genes that are also direct BldD-targets, including *bldM, bldN*, and *whiG* (24). MtrA acts as both activator and repressor in other actinomycetes, but how it impacts *bld* and *whi* gene expression remains to be addressed in *S. venezuelae*. Another example is the MerR-like regulator, BldC, which binds to a number of promoters that are also direct targets of BldD; like MtrA, BldC can have both repressor and activator functions (25).

### Hydrophobic spore coat genes are sensitive to c-di-GMP

The chaplin and rodlin proteins are major components of the hydrophobic sheath that covers the aerial hyphae and spores in *Streptomyces* (26, 27). *S. venezuelae* secretes two long (ChpB and ChpC) and five short (ChpD-H) chaplins, and these proteins are expected to self-assemble into amyloid-like filaments on the cell surface, where they then permit the aerial hyphae to escape the surface tension. As further components of the hydrophobic layer, *S. venezuelae* produces three rodlin proteins (RdlA-C), which are proposed to organize the chaplin filaments into so-called rodlets. Unlike the chaplins, however, the rodlins are dispensable for aerial development and surface hydrophobicity (28). Moreover, when grown on rich medium, *Streptomyces* secrete an additional surfactant peptide, SapB (product of the *ramCSAB* operon) (29).

Expression of genes encoding the different hydrophobic sheath components was significantly affected in the four tested mutants. As shown using RNA-seq, deleting *cdgB* resulted in upregulation of *chpD, chpF, chpG, rdlA, rdlC, ramS* and *ramC* (Fig. 2D). In addition, quantitative RT-PCR (qRT-PCR) data revealed that *chpH* was also upregulated in a *cdgB* mutant (Fig. 3A). Surprisingly, our data showed that in contrast to Δ*cdgB*, all chaplin genes, except *chpB, chpD*, and the three rodlin genes, *rdlA-C*, were downregulated in the *cdgC* mutant (Fig. 2D), despite this strain having the same rapid sporulation phenotype as the *cdgB* mutant. qRT-PCR data confirmed that expression of *chpC, chpE* and *chpH* was 11-fold, 21-fold and 11-fold, respectively, lower in Δ*cdgC* than in wild type (Fig. 3A). We also detected a strong downregulation of the chaplin and rodlin genes in both Δ*rmdA* and Δ*rmdB* strains (Fig. 2D and 3A).

**Figure 3:**
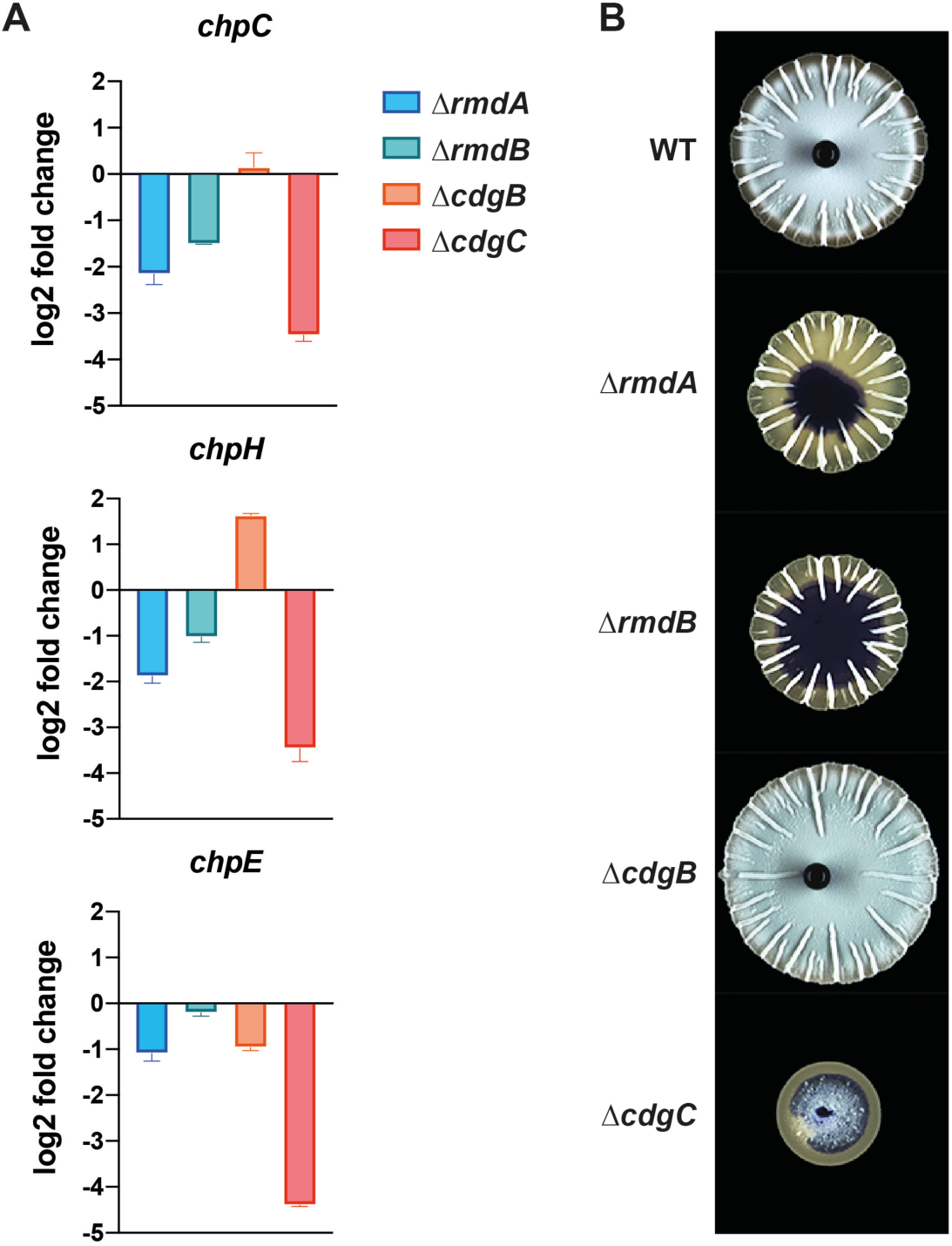
Chaplin expression in the PDE / DGC mutants and properties of their colony surfaces. (A) Strains were grown on MYM plates for 30 h at 30 °C when harvested for RNA isolation that was used for qRT-PCR analysis. The assay was reproduced at least three times with three technical replicates per experiment. Expression values were calculated relative to the accumulation of the constitutively expressed *hrdB* reference mRNA and normalized to the wild type value. log2 change >1/<-1 was considered as significant. Data are presented as mean of technical replicates ± standard deviation (n=3). (B) Twelve µl of 2×10^5^ CFU/µl *S. venezuelae* spores were spotted on MYM agar and incubated for 30 h at 30 °C. Five µl of stained water was pipetted on top of the macrocolonies and images were taken using a binocular camera (Zeiss).

We tested the water repellent properties of the colony surface of the different wild type and mutant strains, and found that wild type and Δ*cdgB* both repelled aqueous solutions (seen as pearl droplet formation on the colony surface), suggesting that they possessed a hydrophobic layer atop their colonies. In contrast, Δ*rmdA*, Δ*rmdB* and Δ*cdgC* colonies were highly hydrophilic, with water droplets immediately dispersing (Fig. 3B). The observed properties associated with these colony surfaces are consistent with expression of *chp* genes in wild type and Δ*cdgB*, and reduced expression of the chaplin genes in Δ*cdgC*, Δ*rmdA* and Δ*rmdB*.

We wondered whether chaplin overexpression could restore the inability of Δ*cdgC*, Δ*rmdA* and Δ*rmdB* to form aerial mycelium. To test this, we introduced *chpB*-*F* and *chpH*, under the control of the constitutive *ermE\** promoter, on the integrative pMS82 vector into each mutant strain. Colony morphology analysis revealed that none of the overexpressed chaplin genes could fully restore aerial mycelium formation to the studied mutants, when overexpressed individually (Fig. S4). Presumably, fine-tuned expression of multiple *chp* genes is needed to overcome this developmental defect (30).

In conclusion, our data revealed that production of the amyloid-forming chaplin and rodlin proteins is controlled by c-di-GMP in *S. venezuelae*. This is reminiscent of many bacteria, in which the synthesis of equivalent extracellular matrix components is activated by c-di-GMP. For example, in *E. coli*, expression of *csgA* and *csgB*, encoding the main components of the amyloid curli fibers, is activated by c-di-GMP (13). However, strikingly, *chp* and *rdl* genes are downregulated upon deletion of the DGC *cdgC*, while deletion of the DGC *cdgB* has the opposite effects, leading to upregulation of these genes. The contrasting expression profile of these genes in the two DGC mutants explains the morphological difference between them. Obviously, lack of a hydrophobic layer means Δ*cdgC* is unable to break the surface tension at the air-agar interface and raise aerial hyphae, so that instead the spores are formed on the upper layer of the substrate mycelium. The downregulation of *chp* and *rdl* genes in Δ*cdgC* is likely a result of *bldN* downregulation in this strain (Fig. 2D and S3A), where *bldN* encodes an ECF sigma factor needed for expression of these genes. *bldN* expression is governed by BldD-(c-di-GMP), while BldN activity is controlled by the membrane-bound anti-sigma factor, RsbN (31). Since CdgC is associated with the membrane via its transmembrane helices, it will be interesting to test whether this enzyme affects *chp* and *rdl* expression through its modulation of RsbN activity.

### Cell division genes are upregulated in the DGC mutants and downregulated in the PDE mutant strains

Our RNA-seq data showed that multiple genes encoding components of the cell division, cell wall synthesis and chromosome segregation machineries, were upregulated upon deletion of *cdgC* (Fig. 2D). Among these targets were *ssgB*, whose product is important for the assembly of FtsZ rings at cell division sites (32); *ssgD*, encoding a protein that appears to be involved in lateral cell wall synthesis; and *ssgE*, whose product was proposed to control the correct timing of spore dissociation (33). In addition, the three *Streptomyces mreB*-like genes (*mreB, vnz35885* and *mbl*) and *mreC* were upregulated in Δ*cdgC* (Fig. 2D). MreB, Mbl and MreC have crucial roles in the synthesis of a thickened spore wall and contribute to resistance of spores to various stresses such as heat, detergents and salt stress (34, 35). The *smeA*-*sffA* operon, which encodes SffA, a putative DNA translocase that participates in chromosome segregation into spores, and the membrane protein SmeA, which localizes SffA to sporulation septa (36), was highly upregulated in Δ*cdgB* and Δ*cdgC* and downregulated in Δ*rmdA* (Fig. 2D).

Differentiation of *Streptomyces* hyphae into spores requires the conserved tubulin-like GTPase FtsZ, which polymerizes into filaments, called Z-rings, close to the membrane and recruits additional cell division proteins (37, 38). Ladder-like array of multiple FtsZ rings define the future sporulation septa. In *S. coelicolor, ftsZ* expression is controlled by three promoters (39); the same organization was observed for the *ftsZ* promoter region in *S. venezuelae* (Fig. 4A). The onset of sporulation coincides with a strong upregulation of *ftsZ* transcription, and this increased expression is crucial for sporulation septation (39). We expected to detect increased *ftsZ* transcript levels in the *cdgB* and *cdgC* mutants that sporulate precociously, but RNA-seq did not reveal significant changes in *ftsZ* expression in any of the mutants. Since the two DGC mutant strains have already formed spores when harvested for RNA-isolation from plates after 30 h of growth, we suspected that harvesting at an earlier time point may have revealed changes in *ftsZ* transcript levels.

**Figure 4:**
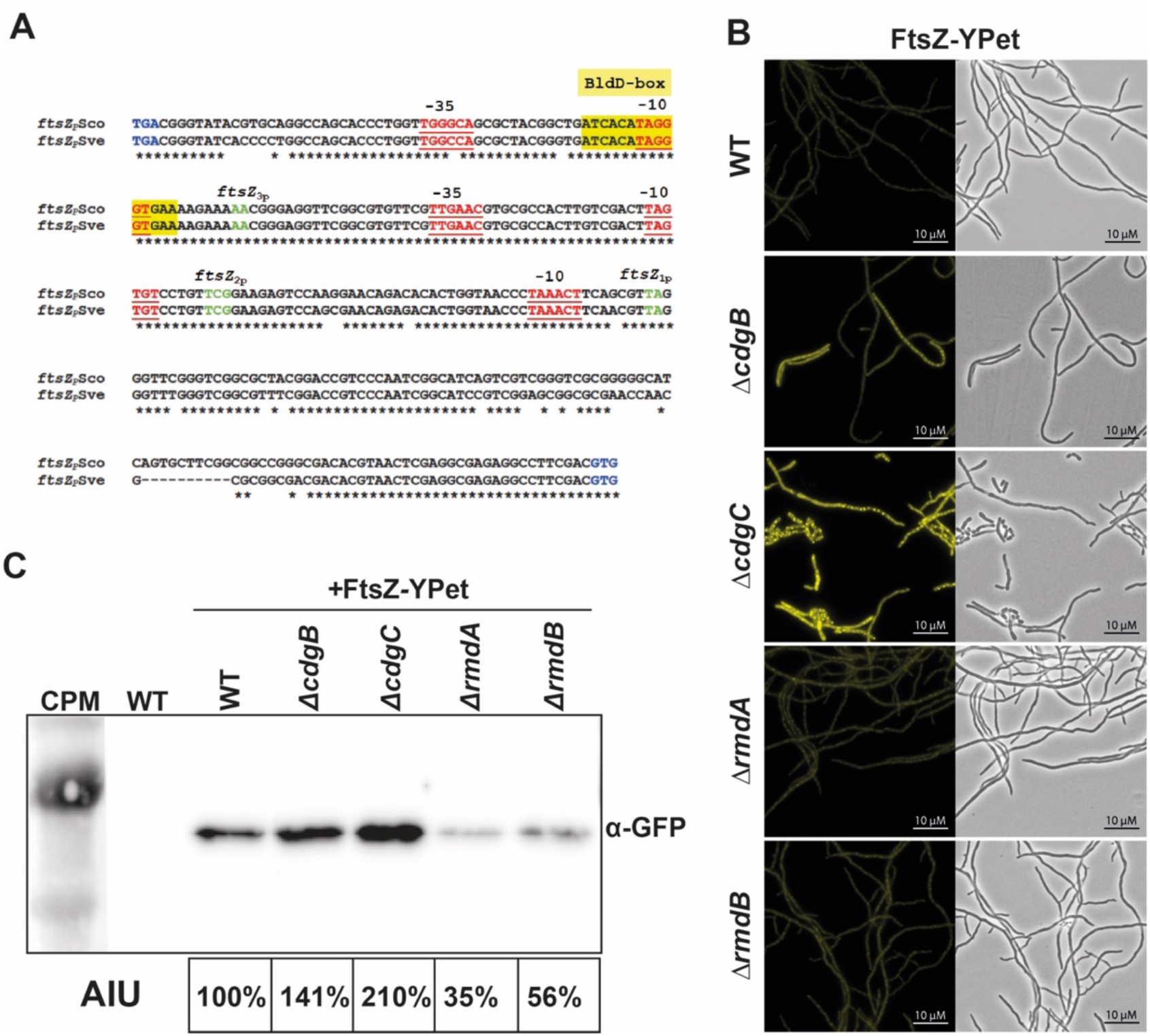
*ftsZ* expression in the *cdgB, cdgC, rmdA* and *rmdB* mutants. (A) *ftsZ* promoter region from *S. coelicolor* (*ftsZ*_P_Sco) and *S. venezuelae* (*ftsZ*_P_Sve). The TGA stop codon from *ftsQ* and the GTG start codon from *ftsZ* are shown in blue. Three *ftsZ* mRNA 5’ends were mapped in the study by Flärdh *et al*., 2000 (39) and are highlighted in green (*ftsZ*_1P_, *ftsZ*_2P_, *ftsZ*_3P_). Putative −10 and −35 promoter regions are underlined and marked in red. BldD-binding site was determined by den Hengst *et al*., 2010 (20) in *S. coelicolor* and is fully conserved in *S. venezuelae* (yellow box). (B) Fluorescence (left) and phase contrast microscopy images (right) showing that FtsZ-YPet is upregulated in Δ*cdgB* and Δ*cdgC* after 12 h of growth in liquid MYM. FtsZ-YPet was expressed from the Φ_BT1_ integration site under control of the native *ftsZ* promoter from the pSS5 vector (40). (C) Immunoblot analysis using anti-GFP antibody for FtsZ-YPet detection. Strains were grown for 12 h in liquid MYM. Fourteen µg of total protein was loaded per lane (see Fig. S5 for loading control). WT free of the FtsZ-YPet fusion was used as negative control. For quantification, arbitrary units (AIU) were determined using ImageQuantTL. CPM: color prestained protein marker (NEB).

Given this, we sought to address *ftsZ* expression in our mutant strains using an alternative approach. We introduced an *ftsZ-ypet* translational fusion, under the control of the native *ftsZ* promoter on the pSS5 plasmid (40), into the Φ_BT1_ phage integration site in the wild type strain, alongside the *cdgB, cdgC, rmdA* and *rmdB* mutants. After 12 h of growth in liquid MYM medium, wild type and the two PDE mutant strains grew vegetatively and only weak FtsZ-YPet signals were detected. In contrast, in the two DGC mutants, the *ftsZ*::*ypet* fusion was highly upregulated, with abundant Z-ring ladders observed, signaling the initiation of sporulation septation. In Δ*cdgC*, single spores were already detectable at this early stage of growth (Fig. 4B). Immunoblot analysis using an anti-GFP antibody confirmed that FtsZ::YPet was most abundant in Δ*cdgC*, and was elevated in Δ*cdgB* relative to the wild type. In contrast, in Δ*rmdA* and Δ*rmdB*, FtsZ::YPet levels were strongly reduced when compared with wild type levels (Fig. 4C).

BldD integrates c-di-GMP signals into *ftsZ* transcriptional control since BldD-(c-di-GMP) directly binds to the *S. venezuelae ftsZ* promoter region, as detected using ChIP-seq analysis (6). The BldD-binding site in the *ftsZ* promoter was defined in *S. coelicolor* (20) and is fully conserved in *S. venezuelae* (Fig. 4A). It is likely that deletion of either *cdgB* or *cdgC* leads to dissociation of the BldD repressor from the *ftsZ* promoter, while deletion of *rmdA* or *rmdB* results in prolonged BldD-(c-di-GMP)-mediated repression. In addition, *ftsZ* expression responds to c-di-GMP changes via σ^WhiG^, which is kept inactive by RsiG-(c-di-GMP) when c-di-GMP levels are high. As demonstrated in *S. coelicolor*, deleting *whiG* or one of the two σ^WhiG^-dependent genes, *whiI* and *whiH*, reduced or eliminated the developmental increase in *ftsZ* transcript levels (39). Altogether, *ftsZ* expression represents a powerful c-di-GMP-sensitive reporter in *Streptomyces*, responding to both, BldD-mediated c-di-GMP-signaling during vegetative growth, and to RsiG-σ^WhiG^-sensed c-di-GMP stimuli during the transition to sporulation.

### Genes encoding second messenger enzymes with altered expression in the DGC / PDE mutants

*In vivo* ChIP-seq analysis identified *cdgA, cdgB, cdgC* and *cdgE* as direct BldD-(c-di-GMP) targets in *S. venezuelae* (6). For *cdgB*, this finding was confirmed biochemically using EMSAs (23), but such confirmation had not been performed for *cdgA, cdgC* and *cdgE*. We systematically tested binding of BldD to promoters of all genes coding for c-di-GMP-metabolizing enzymes in *S. venezuelae* using EMSAs. Our *in vitro* data confirmed that BldD bound in a c-di-GMP-responsive manner to the promoter regions of *cdgA, cdgC* and *cdgE* (Fig. 5A), but we did not detect any protein binding to the promoters of *cdgD, cdgF, rmdA, rmdB* and *hdgAB* (data not shown). BldD binds to a pseudo-palindromic sequence, designated the BldD box; such boxes were located 215, 224 and 59 bp upstream of the translational start codons of *cdgA, cdgC* and *cdgE*, respectively (Fig. 5B). CdgA, CdgB and CdgC are active DGCs (8, 16, 20). We sought to test the DGC activity for CdgE (possessing GAF-GGDEF domains), and found that indeed it too had DGC activity (Fig. 5C). Intriguingly, CdgE activity was subject to product inhibition, since added non-labelled c-di-GMP inhibited conversion of [^32^P]GTP into [^32^P]c-di-GMP (Fig. 5C).

**Figure 5:**
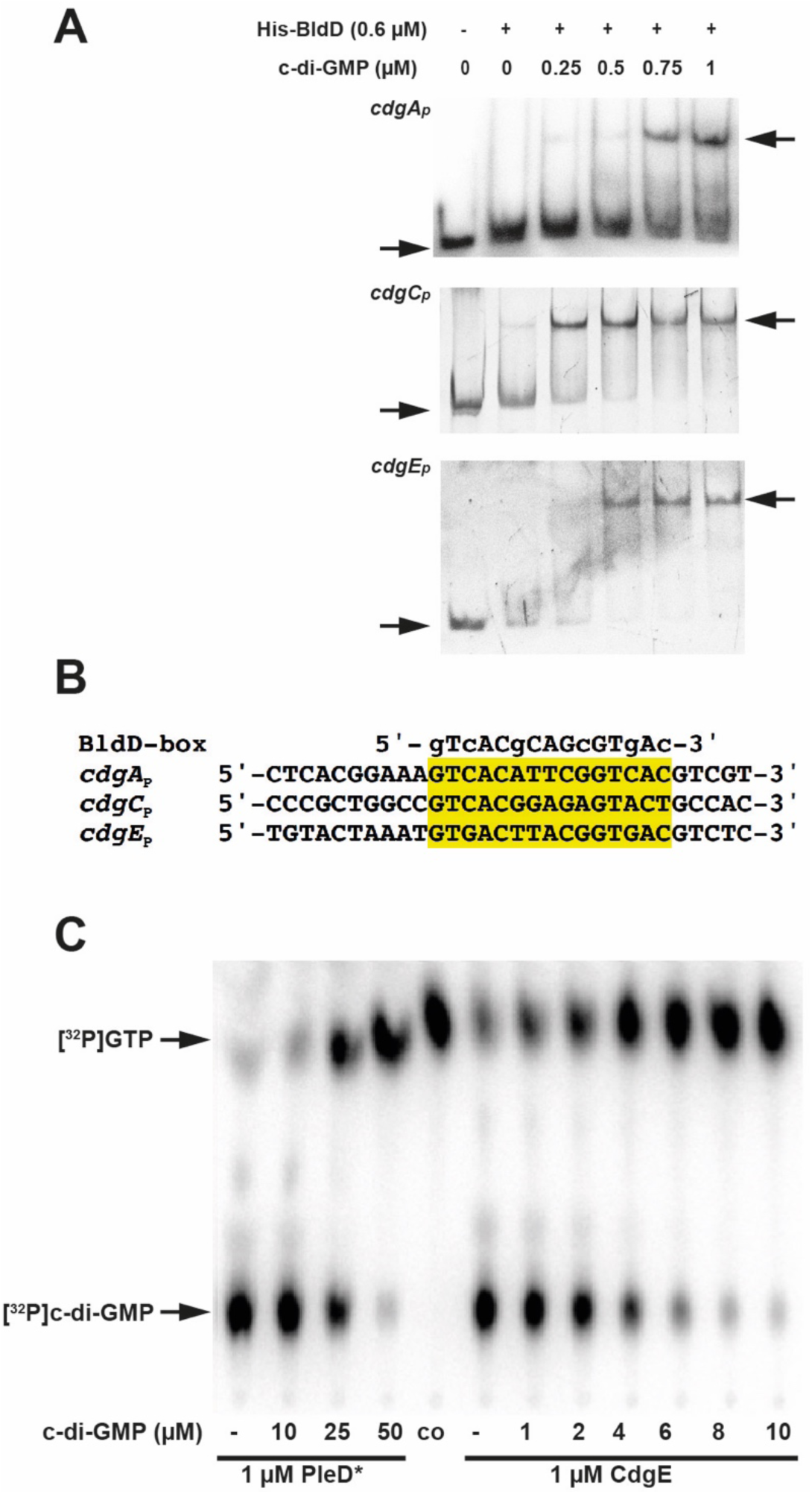
*cdgA, cdgC* and *cdgE* are direct BldD targets. (A) EMSA analysis of BldD binding to *cdgA, cdgC* and *cdgE* promoter DNA ± c-di-GMP (0.25-1 µM). (B) Putative BldD-binding box (yellow) in the promoter regions of *cdgA, cdgC* and *cdgE*. DNA consensus motif bound by BldD was determined by den Hengst *et al*., 2010 (20) and is located 215 bp upstream of the GTG start in *cdgA*_P_, 224 bp upstream of the GTG start in *cdgC*_P_ and 59 bp upstream of the ATG start in *cdgE*_P_. (C) Enzyme assay shows that CdgE is an active DGC and that non-labelled c-di-GMP inhibits CdgE-mediated conversion of [^32^P]GTP into [^32^P]c-di-GMP. The DGC PleD* from *C. crescentus* served as positive control. Co: [^32^P]GTP control.

This regulatory feedback loop comprising BldD as c-di-GMP sensor that controls expression of four active DGCs let us hypothesize that expression of *cdgA, cdgB, cdgC* and *cdgE* may be altered in the analyzed DGC / PDE mutant strains. However, according to RNA-seq, neither transcript abundance of *cdgA*, nor that of *cdgE*, was affected at the tested time point in any of the mutants (Fig. 2D). *cdgC* expression was reduced upon *rmdA* deletion, while *cdgB* transcript levels were lower in Δ*cdgC* than in wild type (Fig. 2D and S3B). Deleting *cdgC* also resulted in downregulation of *rmdA* and upregulation of *cdgF* (Fig. 2D and S3B), which codes for a PAS-PAC-GGDEF-EAL protein that contains 10 predicted transmembrane helices (7).

Transcriptional regulation of c-di-GMP-metabolizing enzymes in *S. venezuelae* is complex and involves the action of multiple global regulators, likely explaining why BldD activity modulation due to changes in c-di-GMP levels in the tested DGC / PDE mutants was not associated with significant transcriptional changes in these genes, at least under the studied conditions. The four direct BldD-(c-di-GMP) targets (*cdgA, cdgB, cdgC* and *cdgE*) are also directly controlled by the response regulator MtrA, which further binds to the promoters of *cdgF* and *rmdB* (24). Moreover, *cdgB* is directly repressed by the transcription factor WhiA, while *cdgE* is directly activated by the MerR-like regulator BldC (41, 42). Such multi-layered transcriptional control of c-di-GMP synthesis and degradation suggests that levels of this molecule are fine-tuned in response to disparate signal transduction cascades.

In addition to genes coding for c-di-GMP-turnover enzymes, we found that *rshA*, encoding a RelA / SpoT homologue containing a conserved HD-domain for hydrolysis of the alarmone (p)ppGpp (7) was downregulated in Δ*cdgC* and Δ*rmdA* (Fig. 2D). In addition, *cya*, encoding a cAMP synthetase was upregulated in Δ*cdgC*, suggesting that CdgC links c-di-GMP-signaling to (p)ppGpp and cAMP metabolism.

### Natural product genes differentially expressed in Δ*cdgB* and Δ*cdgC*

*Streptomyces* spore pigments are frequently aromatic polyketides that are produced by enzymes encoded in the highly conserved *whiE* cluster. In *S. coelicolor*, this cluster comprises an operon of seven genes (*whiE*-ORFI to *whiE*-ORFVII; *sco_5320* – *sco_5314*) and the divergently transcribed gene *whiE*-ORFVIII (*sco_5321*) (43). In *S. venezuelae*, the homologous cluster is similarly organized and encompasses the genes *vnz_33525* to *vnz_33490*. In the *cdgB* and *cdgC* mutants, *whiE*-ORFI to *whiE*-ORFVII genes were up to 12-fold upregulated (Fig. 2D).

Since the *whiE* genes are developmentally regulated and expressed only in spores (43), their upregulation correlates with the morphology of Δ*cdgB* and Δ*cdgC* strains that had already sporulated after 30 h of growth on MYM agar. In contrast, *S. venezuelae* wild type, Δ*rmdA* and Δ*rmdB* were still in the vegetative phase after same incubation period (Fig. S2) and were not expressing the *whiE* genes. *whiE* expression is controlled by the sporulation-specific BldM-WhiI heterodimer (44). Since *whiI* is transcribed in an RsiG-(c-di-GMP)-σ^WhiG^-controlled manner (11), this regulatory circuit is likely responsible for the *whiE* sensitivity to c-di-GMP.

Modulation of c-di-GMP can be an efficient way to manipulate antibiotic production in *Streptomyces* (17). Therefore, we were also interested in identifying antibiotic genes whose expression changed in response to deletion of Δ*cdgB*, Δ*cdgC*, Δ*rmdA* or Δ*rmdB. S. venezuelae* NRRL B-65442 produces the bacteriostatic antibiotic chloramphenicol, a potent inhibitor of bacterial protein biosynthesis. The chloramphenicol biosynthetic gene cluster comprises 17 *cml* genes (*vnz_04400* – *vnz_04480*). These genes were significantly downregulated in both Δ*cdgB* and Δ*cdgC* strains, but were unaffected in Δ*rmdA* and Δ*rmdB* (Fig. 2D). The direct BldD-(c-di-GMP) target gene *bldM* was reported to indirectly repress chloramphenicol genes (45) and may represent a link between c-di-GMP signals and chloramphenicol gene expression.

## Conclusions

The DGCs (CdgB CdgC) and the PDEs (RmdA and RmdB) antagonistically control expression of *ftsZ* via the c-di-GMP-sensors BldD and σ^WhiG^. Upregulation of *ftsZ* together with other cell division genes in the DGC mutants is associated with precocious sporulation, while reduced expression of *ftsZ* in the PDE mutants presumably delays sporulation-specific cell division. Thus, c-di-GMP-responsive expression of cell division genes likely contributes to the decision *when* the spores are formed. In addition, expression of chaplin and rodlin genes – encoding the major components of the hydrophobic sheath that covers the aerial hyphae and spores in *Streptomyces* – is controlled by c-di-GMP. Their expression in combination with the transcriptional profile of cell division genes determines *where* the spores are made: on aerial hyphae or out of substrate mycelium. The c-di-GMP enzymes studied here contribute to balanced combination of cell division components and hydrophobins for coordinated progression of the *Streptomyces* life cycle.

## Material and Methods

### Bacterial strains, plasmids and oligonucleotides

All strains, plasmids and oligonucleotides used in this study are listed in Tables S1 and S2 in the supplemental material. *E. coli* strains were grown in LB medium under aerobic conditions. When required, LB was supplemented with 100 µg/ml ampicillin (Amp), 50 µg/ml kanamycin (Kan), 50 µg/ml apramycin (Apr) or 15 µg/ml chloramphenicol (Cam). For hygromycin B (Hyg) – based selection, nutrient agar (NA; Roth) or LB without NaCl (LBon) were used, to which 16 µg/ml or 22 µg/ml, respectively, Hyg were added. *S. venezuelae* strains (Table S2) were grown aerobically at 30 °C in liquid Maltose-Yeast Extract-Malt Extract (MYM) medium (46) or on MYM agar, both supplemented with trace element solution (47). Liquid cultures were inoculated with spores to a final concentration of 10^6^ colony-forming-units (CFU) per ml. To study development on MYM agar, 12 µl of 2×10^5^ CFU/µl *S. venezuelae* spores were spotted on MYM agar and incubated for the indicated period of time. For hydrophobicity tests, 5 µl of ddH_2_O stained with Coomassie Brilliant Blue G-250 were pipetted on top of the colonies that were grown for 30 h. The resulting macrocolonies were photographed using a binocular (Stemi 2000C, Zeiss) coupled with a camera (AxioCAM ICc 3, Zeiss). Digital images were edited using Photoshop CS6 and Illustrator CS6 software (Adobe).

### Generation of *S. venezuelae* mutant strains

To generate *rmdA*^*ALLEF*^, *rmdA*^*AAA*^ and *rmdB*^*AAA*^ mutations on the SV3-B05 and SV2-B03 cosmid, respectively, recombineering using single-strand oligonucleotides (Table S1) in *E. coli* HME68 was performed as described in (48). Prior to this, the *kan*-resistance cassette of both cosmids was replaced by *apr-oriT* in *E. coli* BW25113/pIJ790. For that, the *apr-oriT* sequence with *neo*-specific extensions was amplified by PCR from pIJ773 (Table S1 and S2). Successful mutagenesis was confirmed by PCR and Sanger sequencing and the confirmed mutant cosmids were transformed into *E. coli* ET12567/pUZ8002 for conjugation into *S. venezuelae*, as described in (22). Conjugation plates were incubated at room temperature overnight, and then overlayed with Apr. Ex-conjugants were re-streaked once on plates containing Apr and nalidixic acid, and then several times on non-selective medium. The desired mutants arising from a double crossing over were screened for Apr-sensitivity followed by PCR to confirm the desired mutations. PCR products comprising the mutagenized regions were sequenced and the resulting strains were named SVJH29 (*rmdA*^*ALLEF*^), SVJH30 (*rmdA*^*AAA*^) and SVJH31 (*rmdB*^AAA^).

### Complementation of Δ*rmdB*

For complementation analysis of Δ*rmdB* with *rmdB*^GGAAF^-FLAG, pIJ10170-*rmdB*^GGAAF^-FLAG was constructed using PCR with pSVJH02 containing *rmdB*-FLAG under the control of the native promoter (8) as a template and the PRJH36 / PRJH37 primer pair (Table S1). The resulting pSVJH03 plasmid was introduced into the phage integration site Φ_BT1_ in the Δ*rmdB* mutant by conjugation and the strain was named SVJH4.

### Immunoblot analysis

For detection of FtsZ-YPet, *S. venezuelae* strains expressing *ftsZ-ypet* controlled by the native *ftsZ* promoter on the pSS5 vector (40) integrated at the Φ_BT1_ phage site, were grown in liquid MYM for 12 h. Two ml were harvested, washed and homogenized in lysis buffer (20 mM Tris, pH 8, 0.5 mM EDTA and cOmplete protease inhibitor cocktail tablets, EDTA-free (Roche)) using a BeadBeater (Biozym; six cycles at 6,00 m/s; 30 s pulse; 60 s interval). Total protein concentration was determined using the Bradford Assay (Roth) and each sample was adjusted to 1 mg/ml. Fourteen µg total protein were loaded per lane and separated on a 12% SDS polyacrylamide gel via electrophoresis and transferred to a polyvinylidene difluoride membrane (PVDF, Roth). For immunodetection, anti-GFP antibody was used and bound primary antibody was detected using anti-rabbit IgG-HRP secondary antibody following visualization with the Clarity™ Western ECL Substrate (BioRad) and subsequent detection in a ECL Chemocam Imager (Intas Pharmaceuticals Limited). For semi-quantitative densitometric evaluation of detected FtsZ-YPet, ImageQuantTL software (GE Healthcare Life Sciences) was used to calculate the amount of pixel per band in equal sized areas indicated as arbitrary intensity units (AIU). Signals were normalized to FtsZ-YPet in wild type that were set to 100%.

### Protein overexpression and purification

Plasmids for overexpression of *cdgE, rmdB* (amino acids 244-704) and *rmdA* (amino acids 164-721) were generated using PCR with oligonucleotides listed in Table S1 and either genomic DNA, pSVJH01 or pSVJH02 (8) as templates. *cdgE* and *rmdB* were cloned into pET15b (Novagen), *rmdA* into pMAL-c2 (NEB). *rmdA* ^*G368G;G369G;D370A;E370A;F371F*^ (*rmdA*^GGAAF^) was created using site directed mutagenesis using the pMAL-c2-*rmdA* plasmid as template. Protein overexpression was induced with IPTG during logarithmic growth of *E. coli* BL21 (DE3) pLysS containing relevant plasmids. 6×His-CdgE and 6×His-RmdB were purified via Ni-NTA chromatography. For MBP-RmdA and MBP-RmdA^GGAAF^ purification, amylose resin (NEB) was applied. For details, please see supplemental material and methods.

### DGC and PDE assay

Enzymatic activity of RmdA, RmdA^GGAAF^, RmdB and CdgE was tested *in vitro* in PDE and DGC assays, respectively, as described in (12, 49) with slight modifications. One µM purified protein in cyclase reaction buffer (25 mM Tris HCl, pH 7.5; 250 mM NaCl with 10 mM MnCl_2_ or MgCl_2_; 5 mM β-mercaptoethanol; 10% glycerol) was incubated with 4.16 nM [^32^P]GTP (Hartmann Analytic GmbH) or 2.08 nM [^32^P]c-di-GMP (Hartmann Analytic GmbH) at 30 °C for 60 min. To stop the reaction, 5 µl 0.5 M EDTA, pH 8 was added to an equal volume of reaction mixture followed by heating to 95 °C for 5 min. In DGC assays, PleD*, a constitutive active DGC from *C. crescentus* (14), was added as positive control. In PDE assays, PdeH from *E. coli* (13) served as a positive control. Samples were separated by thin layer chromatography on Polygram CEL 300 PEI cellulose TLC plates (Macherey– Nagel) incubated in 1:1.5 (v/v) saturated (NH_4_)_2_SO_4_ and 1.5 M KH_2_PO_4_; pH 3.6. After drying, the plates were exposed on a Phosphor Imaging Screen (Fuji) which was then scanned using a Typhoon Scanner FLA 7000 (GE).

### EMSA

Promotor regions of *cdgA* (172 bp), *cdgC* (205 bp) and *cdgE* (121 bp) were amplified by PCR using specific oligonucleotides (Table S1). Twenty ng of DNA was incubated with 0.6 µM His-tagged BldD, 0.5 µg poly[d(I-C)] (Roche) competitor DNA, and increasing concentrations of c-di-GMP. Each sample was supplemented with 2 µl 10× Bandshift buffer (100 mM Tris-HCl, pH 7.5; 100 mM NaCl; 50 mM DTT; 10 mM EDTA; 10 mM MgCl_2_; 50% glycerol) and ddH_2_O to a total volume of 20 µl. Samples were incubated for 20 min at room temperature and loaded onto a 5% polyacrylamide gels prepared with TBE buffer. After separation for 1 hour at 90 V in 0.5 TBE buffer, DNA was visualized by staining with GelRed (Genaxxon) and exposing to UV light.

### RNA isolation, RNA-seq and qRT-PCR

Three *S. venezuelae* macrocolonies that were grown for 30 h at 30°C on MYM agar were pooled and resuspended in 200 µl ice-cold stop solution (5% phenol (pH 4.3) in 98% ethanol) and RNA was isolated using the SV Total RNA Isolation Kit (Promega). After elution, RNA was treated with DNaseI (Turbo DNA-free, Ambion). RNA quantity and quality were analyzed using NanoDrop 2000 (Thermo Scientific) and Bioanalyzer 2100 (Agilent). qRT-PCR was performed using the SensiFAST SYBR No-ROX One-Step Kit (Bioline) and primers listed in Table S1. The RNA-seq libraries were prepared and sequenced in the Illumina NextSeq system by vertis Biotechnologie AG, generating 75 bp single-end reads. The adapter sequences were trimmed from the single-end fastq files using Cutadapt (version 1.18), and low-quality reads were removed.

### Data analysis

Reads were aligned to the *Streptomyces venezuelae* strain NRRL B-65442 genome (accession no. CP018074) using Bowtie 2, with one mismatch allowed. Samtools (version 1.4.1) was used for downstream coverage calculation. The number of reads per gene was obtained using featureCounts (version 1.5.0-p1). The aligned reads were normalized per kilobase per million (RPKM). Differentially transcribed genes were identified using DESeq2 package in R using P-values <0.05 and log2 fold-change <-1 for (downregulated genes) or >1 (for upregulated genes) as significance thresholds. To generate a heat map of differentially expressed genes, we first grouped the targets into selected functional groups. Then we plotted the RPKM normalized values of those genes if they were differentially transcribed in at least one of the *cdgB, cdgC, rmdA* or *rmdB* mutants, using seaborn (version 0.9.0) in Python. To generate Venn diagrams for all the differentially transcribed genes, we used the Venn library (version 0.1.3) in Python. Sequencing data were deposited to the NCBI SRA site under the bioproject accession ID PRJNA608930.

### Phase-contrast and fluorescence microscopy

Before imaging, samples taken from *S. venezuelae* liquid cultures were washed twice in 1× PBS and 5 µl were pipetted on a thin agarose pad on a microscopy slide. Cells were imaged using the Zeiss Axio Observer Z.1 inverted epifluorescence microscope at 100× magnification and the Axiocam 506 mono. Digital images were organized using ADOBE Photoshop software.

## Acknowledgements

We thank Andreas Latoscha and Mirka E. Wörmann for comments on the manuscript. Research in Natalia Tschowri’s lab is funded by the DFG Emmy Noether-Program (TS 325/1-1) and the DFG Priority Program SPP 1879 (TS 325/2-1 and TS 325/2-2), and in Marie Elliot’s lab by the Natural Sciences and Engineering Council of Canada’s Discovery Grant program (RGPIN-2015-04681).

## Author contributions

N.T. designed the study. Experiments were designed, performed and analyzed by J.H., S.A.N, M.M.A-B, S.L and N.T. Scientific consultation by M.A.E. The paper was written by N.T with input from the other authors.

## References

1. Jenal U, Reinders A, Lori C. 2017. Cyclic di-GMP: second messenger extraordinaire. Nat Rev Microbiol 15:271–284.

2. Hengge R. 2009. Principles of c-di-GMP signalling in bacteria. Nat Rev Microbiol 7:263–73.

3. Sarenko O, Klauck G, Wilke FM, Pfiffer V, Richter AM, Herbst S, Kaever V, Hengge R. 2017. More than Enzymes That Make or Break Cyclic Di-GMP-Local Signaling in the Interactome of GGDEF/EAL Domain Proteins of Escherichia coli. MBio 8.

4. Lim B, Beyhan S, Meir J, Yildiz FH. 2006. Cyclic-diGMP signal transduction systems In *Vibrio cholerae*: modulation of rugosity and biofilm formation. Mol Microbiol 60:331–48.

5. Römling U, Galperin MY, Gomelsky M. 2013. Cyclic di-GMP: the first 25 years of a universal bacterial second messenger. Microbiol Mol Biol Rev 77:1–52.

6. Tschowri N, Schumacher MA, Schlimpert S, Chinnam NB, Findlay KC, Brennan RG, Buttner MJ. 2014. Tetrameric c-di-GMP mediates effective transcription factor dimerization to control *Streptomyces* development. Cell 158:1136–47.

7. Latoscha A, Wormann ME, Tschowri N. 2019. Nucleotide second messengers In *Streptomyces*. Microbiology 165:1153–1165.

8. Al-Bassam MM, Haist J, Neumann SA, Lindenberg S, Tschowri N. 2018. Expression Patterns, Genomic Conservation and Input Into Developmental Regulation of the GGDEF/EAL/HD-GYP Domain Proteins In *Streptomyces*. Front Microbiol 9:2524.

9. Hull TD, Ryu MH, Sullivan MJ, Johnson RC, Klena NT, Geiger RM, Gomelsky M, Bennett JA. 2012. Cyclic di-GMP phosphodiesterases RmdA and RmdB are involved in regulating colony morphology and development In *Streptomyces coelicolor*. J Bacteriol 194:4642–51.

10. Bush MJ, Tschowri N, Schlimpert S, Flärdh K, Buttner MJ. 2015. c-di-GMP signalling and the regulation of developmental transitions in streptomycetes. Nat Rev Microbiol 13:749–60.

11. Gallagher KA, Schumacher MA, Bush MJ, Bibb MJ, Chandra G, Holmes NA, Zeng W, Henderson M, Zhang H, Findlay KC, Brennan RG, Buttner MJ. 2020. c-di-GMP Arms an Anti-sigma to Control Progression of Multicellular Differentiation In *Streptomyces*. Mol Cell 77:586–599 e6.

12. Christen M, Christen B, Folcher M, Schauerte A, Jenal U. 2005. Identification and characterization of a cyclic di-GMP-specific phosphodiesterase and its allosteric control by GTP. J Biol Chem 280:30829–37.

13. Pesavento C, Becker G, Sommerfeldt N, Possling A, Tschowri N, Mehlis A, Hengge R. 2008. Inverse regulatory coordination of motility and curli-mediated adhesion In *Escherichia coli*. Genes Dev 22:2434–46.

14. Paul R, Weiser S, Amiot NC, Chan C, Schirmer T, Giese B, Jenal U. 2004. Cell cycle-dependent dynamic localization of a bacterial response regulator with a novel diguanylate cyclase output domain. Genes Dev 18:715–27.

15. Lindenberg S, Klauck G, Pesavento C, Klauck E, Hengge R. 2013. The EAL domain protein YciR acts as a trigger enzyme in a c-di-GMP signalling cascade in E. coli biofilm control. EMBO J 32:2001–14.

16. Tran NT, Den Hengst CD, Gomez-Escribano JP, Buttner MJ. 2011. Identification and characterization of CdgB, a diguanylate cyclase involved in developmental processes In *Streptomyces coelicolor*. J Bacteriol 193:3100–8.

17. Makitrynskyy R, Tsypik O, Nuzzo D, Paululat T, Zechel DL, Bechthold A. 2020. Secondary nucleotide messenger c-di-GMP exerts a global control on natural product biosynthesis in streptomycetes. Nucleic Acids Res 48:1583–1598.

18. Yan H, Lu X, Sun D, Zhuang S, Chen Q, Chen Z, Li J, Wen Y. 2020. BldD, a master developmental repressor, activates antibiotic production in two *Streptomyces* species. Mol Microbiol 113:123–142.

19. Weber H, Polen T, Heuveling J, Wendisch VF, Hengge R. 2005. Genome-wide analysis of the general stress response network In *Escherichia coli*: σ^S^-dependent genes, promoters, and sigma factor selectivity. J Bacteriol 187:1591–603.

20. den Hengst CD, Tran NT, Bibb MJ, Chandra G, Leskiw BK, Buttner MJ. 2010. Genes essential for morphological development and antibiotic production In *Streptomyces coelicolor* are targets of BldD during vegetative growth. Mol Microbiol 78:361–79.

21. Elliot MA, Bibb MJ, Buttner MJ, Leskiw BK. 2001. BldD is a direct regulator of key developmental genes In *Streptomyces coelicolor* A3(2). Mol Microbiol 40:257–69.

22. Bibb MJ, Domonkos A, Chandra G, Buttner MJ. 2012. Expression of the chaplin and rodlin hydrophobic sheath proteins In *Streptomyces venezuelae* is controlled by sigma(BldN) and a cognate anti-sigma factor, RsbN. Mol Microbiol 84:1033–49.

23. Schumacher MA, Zeng W, Findlay KC, Buttner MJ, Brennan RG, Tschowri N. 2017. The *Streptomyces* master regulator BldD binds c-di-GMP sequentially to create a functional BldD2-(c-di-GMP)4 complex. Nucleic Acids Res 45:6923–6933.

24. Som NF, Heine D, Holmes NA, Munnoch JT, Chandra G, Seipke RF, Hoskisson PA, Wilkinson B, Hutchings MI. 2017. The Conserved Actinobacterial Two-Component System MtrAB Coordinates Chloramphenicol Production with Sporulation In *Streptomyces venezuelae* NRRL B-65442. Front Microbiol 8:1145.

25. Schumacher MA, den Hengst CD, Bush MJ, L. TBK, Tran NT, Chandra G, Zeng W, Travis B, Brennan RG, Buttner MJ. 2018. The MerR-like protein BldC binds DNA direct repeats as cooperative multimers to regulate *Streptomyces* development. Nat Commun 9:1139.

26. Elliot MA, Karoonuthaisiri N, Huang J, Bibb MJ, Cohen SN, Kao CM, Buttner MJ. 2003. The chaplins: a family of hydrophobic cell-surface proteins involved in aerial mycelium formation In *Streptomyces coelicolor*. Genes Dev 17:1727–40.

27. Claessen D, Rink R, de Jong W, Siebring J, de Vreugd P, Boersma FG, Dijkhuizen L, Wosten HA. 2003. A novel class of secreted hydrophobic proteins is involved in aerial hyphae formation In *Streptomyces coelicolor* by forming amyloid-like fibrils. Genes Dev 17:1714–26.

28. Claessen D, Wosten HA, van Keulen G, Faber OG, Alves AM, Meijer WG, Dijkhuizen L. 2002. Two novel homologous proteins of *Streptomyces coelicolor* and *Streptomyces lividans* are involved in the formation of the rodlet layer and mediate attachment to a hydrophobic surface. Mol Microbiol 44:1483–92.

29. Willey J, Santamaria R, Guijarro J, Geistlich M, Losick R. 1991. Extracellular complementation of a developmental mutation implicates a small sporulation protein in aerial mycelium formation by S. coelicolor. Cell 65:641–50.

30. Di Berardo C, Capstick DS, Bibb MJ, Findlay KC, Buttner MJ, Elliot MA. 2008. Function and redundancy of the chaplin cell surface proteins in aerial hypha formation, rodlet assembly, and viability In *Streptomyces coelicolor*. J Bacteriol 190:5879–89.

31. Schumacher MA, Bush MJ, Bibb MJ, Ramos-Leon F, Chandra G, Zeng W, Buttner MJ. 2018. The crystal structure of the RsbN-sigmaBldN complex from *Streptomyces venezuelae* defines a new structural class of anti-sigma factor. Nucleic Acids Res 46:7467–7468.

32. Willemse J, Borst JW, de Waal E, Bisseling T, van Wezel GP. 2011. Positive control of cell division: FtsZ is recruited by SsgB during sporulation of *Streptomyces*. Genes Dev 25:89–99.

33. Noens EE, Mersinias V, Traag BA, Smith CP, Koerten HK, van Wezel GP. 2005. SsgA-like proteins determine the fate of peptidoglycan during sporulation of *Streptomyces coelicolor*. Mol Microbiol 58:929–44.

34. Heichlinger A, Ammelburg M, Kleinschnitz EM, Latus A, Maldener I, Flärdh K, Wohlleben W, Muth G. 2011. The MreB-like protein Mbl of *Streptomyces coelicolor* A3(2) depends on MreB for proper localization and contributes to spore wall synthesis. J Bacteriol 193:1533–42.

35. Kleinschnitz EM, Heichlinger A, Schirner K, Winkler J, Latus A, Maldener I, Wohlleben W, Muth G. 2011. Proteins encoded by the mre gene cluster In *Streptomyces coelicolor* A3(2) cooperate in spore wall synthesis. Mol Microbiol 79:1367–79.

36. Ausmees N, Wahlstedt H, Bagchi S, Elliot MA, Buttner MJ, Flärdh K. 2007. SmeA, a small membrane protein with multiple functions In *Streptomyces* sporulation including targeting of a SpoIIIE/FtsK-like protein to cell division septa. Mol Microbiol 65:1458–73.

37. Haeusser DP, Margolin W. 2016. Splitsville: structural and functional insights into the dynamic bacterial Z ring. Nat Rev Microbiol 14:305–19.

38. Jakimowicz D, van Wezel GP. 2012. Cell division and DNA segregation In *Streptomyces*: how to build a septum in the middle of nowhere? Mol Microbiol 85:393–404.

39. Flärdh K, Leibovitz E, Buttner MJ, Chater KF. 2000. Generation of a non-sporulating strain of *Streptomyces coelicolor* A3(2) by the manipulation of a developmentally controlled *ftsZ* promoter. Mol Microbiol 38:737–49.

40. Schlimpert S, Wasserstrom S, Chandra G, Bibb MJ, Findlay KC, Flärdh K, Buttner MJ. 2017. Two dynamin-like proteins stabilize FtsZ rings during *Streptomyces* sporulation. Proc Natl Acad Sci U S A 114:E6176–E6183.

41. Bush MJ, Bibb MJ, Chandra G, Findlay KC, Buttner MJ. 2013. Genes Required for Aerial Growth, Cell Division, and Chromosome Segregation Are Targets of WhiA before Sporulation In *Streptomyces venezuelae*. MBio 4.

42. Bush MJ, Chandra G, Al-Bassam MM, Findlay KC, Buttner MJ. 2019. BldC Delays Entry into Development To Produce a Sustained Period of Vegetative Growth In *Streptomyces venezuelae*. MBio 10.

43. Kelemen GH, Brian P, Flärdh K, Chamberlin L, Chater KF, Buttner MJ. 1998. Developmental regulation of transcription of *whiE*, a locus specifying the polyketide spore pigment In *Streptomyces coelicolor* A3 2. J Bacteriol 180:2515–21.

44. Al-Bassam MM, Bibb MJ, Bush MJ, Chandra G, Buttner MJ. 2014. Response regulator heterodimer formation controls a key stage In *Streptomyces* development. PLoS Genet 10:e1004554.

45. Fernandez-Martinez LT, Borsetto C, Gomez-Escribano JP, Bibb MJ, Al-Bassam MM, Chandra G, Bibb MJ. 2014. New insights into chloramphenicol biosynthesis In *Streptomyces venezuelae* ATCC 10712. Antimicrob Agents Chemother 58:7441–50.

46. Stuttard C. 1982. Temperate phages of *Streptomyces venezuelae*: lysogeny and host specificity shown by phages SV1 and SV2. Microbiology 128:115–121.

47. Kieser T, Bibb MJ, Buttner MJ, Chater KF, Hopwood DA. 2000. Practical *Streptomyces* Genetics. The John Innes Foundation, Norwich.

48. Feeney MA, Chandra G, Findlay KC, Paget MSB, Buttner MJ. 2017. Translational Control of the SigR-Directed Oxidative Stress Response In *Streptomyces* via IF3-Mediated Repression of a Noncanonical GTC Start Codon. mBio 8.

49. Weber H, Pesavento C, Possling A, Tischendorf G, Hengge R. 2006. Cyclic-di-GMP-mediated signalling within the σ^S^ network of *Escherichia coli*. Mol Microbiol 62:1014–34.

