## Supplementary Information for "Specialized and shared functions of diguanylate cyclases and phosphodiesterases in *Streptomyces* development"

**This PDF file includes:**

- Extended Experimental Procedures
- Table S1
- Table S2
- Figures S1 to S5
- Supplementary References

### EXTENDED EXPERIMENTAL PROCEDURES

#### Protein overexpression and purification

For protein overexpression from pET15b or pMAL-c2 vector, respectively, *E. coli* BL21 (DE3) pLysS cells were grown in LB supplemented with Amp and Cam (and 2 g/l glucose in the case of pMAL-c2 constructs). Cultures were grown at 37 °C and protein overexpression was induced with a final concentration of 0.3 mM IPTG at $OD_{578}$ of 0.5. Additionally, all cultures were supplemented with either 0.35 mM $MgCl_2$ or $MnCl_2$ (1). Proteins were overexpressed for 4 h at 30 °C or for 20 h at 20 °C. Cells were harvested by centrifugation at 7000 rpm for 15 min. Cell pellets were resuspended in lysis buffer (see below) and homogenized using a Digi-F-Press (G. Heinemann).

Strains expressing 6×His-RmdB, 6×His-RmdB$^{EAL}$ and 6×His-CdgE were lysed in lysis buffer (50 mM Tris HCl, pH 8; 300 mM NaCl; 10% glycerol, 20 mM imidazole; 1 mM ß-mercaptoethanol; 0.1 % Triton X-100) supplemented with cOmplete protease inhibitor cocktail tablets, EDTA-free (Roche). After centrifugation for 45 min at 16.000 rpm, 4 °C, supernatants were combined with 0.5-1 ml 50% Ni-NTA SuperFlow and transferred to a 20 ml column for washing with buffer containing: 50 mM Tris HCl, pH 8; 300 mM NaCl; 10% glycerol, 50 mM imidazole; 1 mM ß-mercaptoethanol; 0.1 % Triton X-100. Proteins were eluted with elution buffer (50 mM Tris HCl, pH 8; 300 mM NaCl; 10% glycerol; 250 mM imidazole; 1 mM ß-mercaptoethanol; 5 mM $MnCl_2$ or $MgCl_2$).

Cells with overexpressed MBP-RmdA and MBP-RmdA$^{GGAAF}$ were lysed in column buffer (50 mM Tris-HCl, pH 7.5; 300 mM NaCl; 10 mM $MnCl_2$; 1 mM ß-mercaptoethanol; 10% glycerol) supplemented with cOmplete protease inhibitor cocktail tablets, EDTA-free (Roche). Cell lysates were centrifuged for 45 min at 16000 rpm, 4 °C and supernatants were incubated with 0.5-1 ml amylose resin (NEB) for 1 h at 4 °C. Afterwards the suspensions were transferred onto a 20 ml column and washed with column buffer. MBP-tagged proteins were eluted using column buffer supplemented with 10 mM maltose.

Purified proteins for usage in enzyme assays were dialyzed twice in cyclase reaction buffer (CRB) (25 mM Tris HCl, pH 7.5; 250 mM NaCl; 10 mM $MnCl_2$ or $MgCl_2$, 5 mM β-mercaptoethanol, 10% glycerol), modified from (2) and stored at -20 °C. The phosphodiesterase PdeH used as positive control in PDE assays was overexpressed and isolated as described in Pesavento *et al.*, 2008 (3). The diguanylate cyclase PleD* which was used as positive control in DGC assays, was overexpressed and purified as described in Paul *et al.,* 2004 (4). For EMSA, His-BldD was overexpressed from pIJ10663 at 37°C and purified using Ni-NTA chromatography exactly as in the study by Tschowri *et al.*, 2014 (5).

**RNA isolation for RNA sequencing and qRT-PCR**

Cell material from three *S. venezuelae* macrocolonies grown on MYM-agar was pooled and resuspended in 200 µl ice-cold stop solution (5% phenol (pH 4.3; Roth) solved in 98% ethanol). After centrifugation, the supernatant was removed and cell pellets were stored at -80 °C. For RNA isolation, pellets were resuspended in 700 µl RLA Buffer (Promega) and then split into two 350 µl fractions. After the addition of 352 µl phenol (Roth, RNA, pH 4.3) and 88 µl chloroform-isoamylalcohol (24:1, AppliChem) to one of the fractions, samples were transferred onto tubes with lysing matrix B (MP Biomedicals). Homogenization was performed using a FastPrep-24™ 5G Instrument (MP Biomedicals). Five pulses of 30 s of intensity 6.0 m/s were applied with cooling for 1 min between pulses. Suspensions were centrifuged for 15 min at 4 °C at 14000 rpm and supernatants were transferred into a new reaction tube. After repetition of this step, 690 µl of RNA Dilution Buffer (Promega) was added to 350 µl of lysate and the reaction was incubated at 70 °C for 3 min. Upon addition of

395 µl 98% ethanol, the samples were treated according to the instructions provided in the SV Total RNA Isolation Kit manual (Promega). After elution with 100 µl RNase free water (Promega), the RNA samples were treated by an additional DNase I treatment (Turbo DNA-free, Ambion) according to the instructions in the manual. RNA quantity and quality were analyzed using NanoDrop 2000 (Thermo Scientific) and Bioanalyzer 2100 (Agilent) measurements.

**qRT-PCR**

qRT-PCR was performed using the SensiFAST SYBR No-ROX One-Step Kit (Bioline). Three technical replicates of each gene were used in every run. RNA samples were adjusted to a concentration of 10 µg/µl and 2 µl were applied in each run. Specific qRT-PCR primers (Table S1, final concentration 250 nM) were used to amplify the target genes (*bldM, bldN, chpC, chpE, chpH, cdgB, cdgC, cdgF* and *rmdA*) and the reference gene *hrdB*. For normalization, primer efficiency was tested by generating a standard curve using chromosomal DNA. Additionally, melting curve analysis was performed to confirm the production of a specific single product from each primer pair. qRT-PCR was run on a CFX Connect real time system (Bio-Rad) using sealed 96-well PCR plates (BioRad). PCR products were amplified according to the following protocol: 45 °C 20 min, 95 °C 5 min, then 35 cycles at 95 °C 10 s, 60 °C 20 s and 72 °C 10 s. The melting curves were generated from 65 to 95 °C with 0.5 °C increments. The experiments were repeated three times independently. The resulting mean starting quantity (SQ) values were averaged and the fold difference was calculated by the standard curve method using following formulas:

$$Fold\ difference = \frac{(E_{\ target})^{\Delta Ct\ target}}{(E_{\ normalizer})^{\Delta Ct\ normalizer}}$$

$$E = efficiency\ from\ standard\ curve\ E = 10^{-\frac{1}{slope}}$$

$$\Delta Ct_{\ target} = Ct_{\ GOI}\ \boldsymbol{c} - Ct_{\ GOI}\ \boldsymbol{s}$$

$$\Delta Ct_{\ normalizer} = Ct_{\ norm}\ \boldsymbol{c} - Ct_{\ norm}\ \boldsymbol{s}$$

Fold difference values were then converted into log2 change fold.

**Table S1. Oligonucleotides used in this study**

| Oligonucleotides | Sequence |
|---|---|
| **Oligonucleotides used for amplification of the *oriT-apr* cassette with *neo*-specific extensions for replacement of the kan$^{R}$ cassette on cosmid SV2-B03 and SV3-B05 and for verification of the exchange** | |
| neo_H1-P1-fw | AGATCTGATCAAGAGACAGGATGAGGATCGTTTCGCatgATTCCGGGG ATCCGTCGACC |
| neo_H2-P2-rev | TCGCTTGGTCGGTCATTTCGAACCCCAGAGTCCCGCtcaTGTAGGCTGG AGCTGCTTC |
| neo_test-fw | GTTTTATGGACAGCAAGCG |
| neo_test-rev | GAATCGAAATCTCGTGATG |
| **Oligonucleotides used for chromosomal *rmdA*$^{ALLEF}$ point mutation, PCR verification and sequencing** | |
| PRJH17 | ACCGGCTCCGGCGAGATGGTCGCCCGGCTCgcactgctgGAGTTCGTCGC CCTCACCACCGGCGCCGCCG |
| Sequence primer rmdA_ALLEF | TGCTGCACCTGCGCCTGCGG |
| Test primer rmdA_ALLEF_fw | GTCGCCCGGCTCGCACTGCTG |
| Test primer rmdA_ALLEF_rw_new | TCAGACGGCGTCCGCCAGCAG |
| **Oligonucleotides used for chromosomal *rmdA*$^{AAA}$ point mutation, PCR verification and sequencing** | |
| PRJH18 | GCACGCCGTGCTGCGGGTGGCACCACCGGACGGCGGCGGCGGCGCC GTGCACACTGCCGTCGTCGAGGTG |
| Sequence primer rmdA_AAA | CTGACCGTCCGCGGCAGCC |
| Test primer rmdA_AAA_fw_new | CGGCGCCGCCGCCGCC |
| Test primer rmdA_AAA_rw_new | TCAGACGGCGTCCGCCAGCAG |
| **Oligonucleotides used for chromosomal *rmdB*$^{AAA}$ point mutation, PCR verification and sequencing** | |

| | |
|---|---|
| PRJH19 | GGTCCAGTTCGACGGCCAGGTCGCCGGCCTCGCCGCGGCCGTCCGCT GGGTCCACCCCGAGCGCGGCAAG |
| Sequence primer rmdB_AAA | CTCCACCACCAGCGCCCAGC |
| Test primer rmdB_AAA_fw_new | CCGGCCTCGCCGCGGCC |
| Test primer rmdB_AAA_rw_new | CTACTGGACGACCTGGCCGGAG |
| **Oligonucleotides used for generation of pET15b-*rmdB* overexpression construct** | |
| PRJH11 | ATACATATGTCCACCCTGTGGATCGC |
| PRJH12 | ATAGGATCCCTACTGGACGACCTGGC |
| **Oligonucleotides used for generation of pET15b-*cdgE* overexpression construct** | |
| 4502-BamHI-pET15b-rev | CTCCTCGGATCCTCACCCGCCCCCGTCCCCG |
| 4502-NdeI-pET15b-fw | GGTGGTCATATGGGTGAGGACGTACGGC |
| **Oligonucleotides used for generation of pET15b-*rmdB*$^{EAL}$ overexpression construct** | |
| PRJH13 | ATACATATGCGCGACTCCAACACCC |
| PRJH12 | ATAGGATCCCTACTGGACGACCTGGC |
| **Oligonucleotides used for generation of pMAL-c2*rmdA* overexpression construct** | |
| PRJH15 | TATATGAATTCAGCCGCTTCCGCGCGGTCTT |
| PRJH16 | TATAAAGCTTGTCAGACGGCGTCCGCCAGCA |
| **Oligonucleotides used for generation of pMAL-c2*rmdA*$^{GGAAF}$ overexpression construct (construction by site-directed mutagenesis based on pMAL-c2*rmdA*)** | |
| PRJH26 | CCCGGCTCGGCGGCGcCGccTTCGTCGCCCTCACC |
| PRJH27 | CGACCATCTCGCCGGAGCCGG |
| **Oligonucleotides used for generation of pMS82$_{permE*}$-*chpC*, -*chpE* and -*chpH* overexpression construct (via integration of the gene sequence into the *ermE** promotor-containing pMC500 vector and subcloning of the promotor + gene into pMS82 vector)** | |

| | |
|---|---|
| ChpC-oe-n-fwd | GACGAGATATCGTGGCACTCTTTCGAAGCCGA |
| ChpC-oe-rev | GACGGATCCCTGTGCGTGGTGCGTG |
| ChpE-oe-fwd | GACGAGAATTCGTGAAGAACCTCAAGAAGGC |
| ChpE-oe-rev | GACGGATCCGGGTGTCAGCCGTTGA |
| ChpH-oe-fwd | GACGAGAATTCATGATCAAGAAGGTCGTCGC |
| ChpH-oe-rev | GACGGATCCGCGAGGTTCAACGTCA |
| **Oligonucleotides used for generation of p3xFLAG-*rmdB*$^{AAL}$ (construction by site-directed mutagenesis based on p3xFLAG-*rmdB*)** | |
| PRJH36 | CCCGGCTCGGCGGCGCCGCCTTCGCCGTCCTGCTG |
| PRJH37 | CGGCCTCGGCGCCGCGGGG |
| **Oligonucleotides used for qRT-PCR** | |
| hrdBqRT_F1 | TGTTCTGCGCAGCCTCAATC |
| hrdBqRT_R1 | CTCTTCGCTGCGACGCTCTT |
| BldM_fw | ATGACATCCGTTCTCGTCTGc |
| BldM_rw | AGGACTTCCTCGCCGTTGG |
| BldN_fw | ACTTCAAGTCCAGCCGGTTC |
| BldN_rw | TTGGAGAGGGACTCCAGGAC |
| chpC_qRT-PCR fw | AACGGTGCGGCCATGAATTC |
| chpC_qRT-PCR rev | CGACGACATTGACCGTGTTTC |
| chpE_qRT-PCR fw | CGTCCCCGTGAACGTCTC |
| chpE_qRT-PCR rev | CCGTTGAGGGCGTGGTTG |
| chpH_qRT-PCR fw | AAGAAGGTCGTCGCTGCTG |
| chpH_qRT-PCR rev | GGACGACATTGCCCGAGAG |
| cdgB_qRT-PCR fw | AGCTGCTCGGCGTCATATC |
| cdgB_qRT-PCR rev | GCACGCTGCATGTTGGAG |

| | |
|---|---|
| cdgC_qRT-PCR fw | AACCACTTCAGGTCACTCGTG |
| cdgC_qRT-PCR rev | ATGAGCGAGGCCAGTTCGG |
| cdgF_qRT-PCR fw | ACCGAACAGCGGAAACTGGAA |
| cdgF_qRT-PCR rev | ATGCAGGTCAGGGTGGATTC |
| rmdA-qRT-PCR_fw | GAGGTCAACGAGACCCTCAC |
| rmdA-qRT-PCR_rw | TCGTACAGCTTCCAGACGTG |
| **Oligonucleotides used for amplification of *cdgA* promoter for EMSA** | |
| pcdgA-fw | CGCATGTTTCCGCTGCC |
| pcdgA-rev | GTCCGGCACTGTGAGGCC |
| **Oligonucleotides used for used for amplification of *cdgC* promoter for EMSA** | |
| 5187-P-Flag-fw | CCTCACCACCACCTGACC |
| 5187_emsa_rev_205 | CTCGGAATCTGATGTCCC |
| **Oligonucleotides used for used for amplification of *cdgE* promoter for EMSA** | |
| 4602_emsa_fw | CAACGAGAAGCGGAAGCC |
| 4602_emsa_rev_121 | CCATTCCTCCAGCTTAGG |

### Table S2. Strains, plasmids and cosmids used in this study

| **Strains** | **Genotype or comments** | **Source or reference** |
|---|---|---|
| ***S. venezuelae*** | | |
| NRRL B-65442 | Wild type | (NCBI Reference Sequence: NZ_CP018074.1) |
| SVNT20 | *cdgC::apr*; Apr$^{R}$ | (6) |
| SVNT23 | *cdgB::apr*; Apr$^{R}$ | (6) |
| SVNT26 | *rmdB::apr*; Apr$^{R}$ | (6) |
| SVNT27 | *rmdA::apr*; Apr$^{R}$ | (6) |

| | | |
|---|---|---|
| SVNT39 | *rmdB::apr; attB*$_{\Phi BT1}$::p3xFLAG-*rmdB*; Apr$^{R}$, Hyg$^{R}$ | (6) |
| SVJH4 | *rmdB::apr; attB*$_{\Phi BT1}$::p3xFLAG-*rmdB*$^{AAL}$; Apr$^{R}$, Hyg$^{R}$ | This study |
| SVJH15 | *rmdA::apr*; *attB*$_{\Phi BT1}$::pMS82, Apr$^{R}$, Hyg$^{R}$ | This study |
| SVJH16 | *rmdB::apr*; *attB*$_{\Phi BT1}$::pMS82, Apr$^{R}$, Hyg$^{R}$ | This study |
| SVJH17 | *rmdA::apr*; *attB*$_{\Phi BT1}$::pMS82$_{-ermE*}$-*chpB*, Apr$^{R}$, Hyg$^{R}$ | This study |
| SVJH18 | *rmdA::apr*; *attB*$_{\Phi BT1}$:: pMS82$_{-ermE*}$-*chpC*, Apr$^{R}$, Hyg$^{R}$ | This study |
| SVJH19 | *rmdA::apr*; *attB*$_{\Phi BT1}$:: pMS82$_{-ermE*}$-*chpD*, Apr$^{R}$, Hyg$^{R}$ | This study |
| SVJH20 | *rmdA::apr*; *attB*$_{\Phi BT1}$:: pMS82$_{-ermE*}$-*chpE*, Apr$^{R}$, Hyg$^{R}$ | This study |
| SVJH21 | *rmdA::apr*; *attB*$_{\Phi BT1}$:: pMS82$_{-ermE*}$-*chpF*, Apr$^{R}$, Hyg$^{R}$ | This study |
| SVJH22 | *rmdA::apr*; *attB*$_{\Phi BT1}$:: pMS82$_{-ermE*}$-*chpH*, Apr$^{R}$, Hyg$^{R}$ | This study |
| SVJH23 | *rmdB::apr*; *attB*$_{\Phi BT1}$:: pMS82$_{-ermE*}$-*chpB*, Apr$^{R}$, Hyg$^{R}$ | This study |
| SVJH24 | *rmdB::apr; attB*$_{\Phi BT1}$:: pMS82$_{-ermE*}$-*chpC*, Apr$^{R}$, Hyg$^{R}$ | This study |
| SVJH25 | *rmdB::apr*; *attB*$_{\Phi BT1}$:: pMS82$_{-ermE*}$-*chpD*, Apr$^{R}$, Hyg$^{R}$ | This study |
| SVJH26 | *rmdB::apr*; *attB*$_{\Phi BT1}$:: pMS82$_{-ermE*}$-*chpE*, Apr$^{R}$, Hyg$^{R}$ | This study |
| SVJH27 | *rmdB::apr*; *attB*$_{\Phi BT1}$:: pMS82$_{-ermE*}$-*chpF*, Apr$^{R}$, Hyg$^{R}$ | This study |
| SVJH28 | *rmdB::apr*; *attB*$_{\Phi BT1}$:: pMS82$_{-ermE*}$-*chpH*, Apr$^{R}$, Hyg$^{R}$ | This study |
| SVSN29 | *cdgC::apr*; *attB*$_{\Phi BT1}$:: pMS82$_{-ermE*}$-*chpB*, Apr$^{R}$, Hyg$^{R}$ | This study |
| SVSN31 | *cdgC::apr*; *attB*$_{\Phi BT1}$:: pMS82$_{-ermE*}$-*chpC*, Apr$^{R}$, Hyg$^{R}$ | This study |
| SVSN33 | *cdgC::apr*; *attB*$_{\Phi BT1}$:: pMS82$_{-ermE*}$-*chpD*, Apr$^{R}$, Hyg$^{R}$ | This study |
| SVSN35 | *cdgC::apr*; *attB*$_{\Phi BT1}$:: pMS82$_{-ermE*}$-*chpE*, Apr$^{R}$, Hyg$^{R}$ | This study |
| SVSN37 | *cdgC::apr*; *attB*$_{\Phi BT1}$:: pMS82$_{-ermE*}$-*chpF*, Apr$^{R}$, Hyg$^{R}$ | This study |
| SVSN39 | *cdgC::apr*; *attB*$_{\Phi BT1}$:: pMS82$_{-ermE*}$-*chpH*, Apr$^{R}$, Hyg$^{R}$ | This study |
| SVJH29 | *rmdA::rmdA*$^{ALLEF}$ | This study |
| SVJH30 | *rmdA::rmdA*$^{AAA}$ | This study |

| | | |
|---|---|---|
| SVJH31 | *rmdB::rmdB*$^{AAA}$ | This study |
| SVSN62 | *cdgB::apr*; *attB*$_{\Phi BT1}$*::* pIJ10750-P$_{ftsZ}$-*ftsZ-ypet,* Hyg$^{R}$ | This study |
| SVSN63 | *cdgC::apr*; *attB*$_{\Phi BT1}$*::* pIJ10750-P$_{ftsZ}$-*ftsZ-ypet,* Hyg$^{R}$ | This study |
| SVJH32 | *rmdA::apr*; *attB*$_{\Phi BT1}$*::* pIJ10750-P$_{ftsZ}$-*ftsZ-ypet,* Hyg$^{R}$ | This study |
| SVJH33 | *rmdB::apr*; *attB*$_{\Phi BT1}$*::* pIJ10750-P$_{ftsZ}$-*ftsZ-ypet,* Hyg$^{R}$ | This study |
| ***E. coli*** | | |
| W3110 | K-12 derivative; *F-, λ- ,rpoS(Am), rph-1, Inv(rrnD-rrnE)* | (7) |
| W3110 Δ*cya* | W3110 derivative with deleted adenylate cyclase | (8) |
| ET12567/pUZ8002 | *dam*, *dcm*, *hsd*; Kan$^{R}$, Cm$^{R}$ | (9) |
| BW25113/pIJ790 | (Δ(*araD-araB*)*567*, Δ*lacZ4787*(*::rrnB-4*), *lacIp-4000*(lacI$^{Q}$), λ-, *rpoS369*(Am), *rph-1*, Δ(*rhaD-rhaB*)568, *hsdR514*; Cm$^{R}$ | (10) |
| BL21 (DE3) pLysS | F– *ompT hsdS*(rB– mB–) *gal dcm* λ(DE3), Cm$^{R}$ | Promega |
| HME68 | W3110 Δ(*argF-lac*)*U169 galKtyr145UAG mutS<>cat* | (11) |
| **Plasmids** | | |
| pIJ773 | Plasmid template for amplification of the *apr-oriT* cassette for 'Redirect' PCR-targeting; Apr$^{R}$ | (12) |
| pIJ790 | Modified λRED recombination plasmid [*oriR101*] [*repA101*(ts)] *araBp-gam-be-exo*; Cm$^{R}$ | (12) |
| pIJ10770 | pMS82 derivative; Hyg$^{R}$ | (13) |
| pIJ10663 | pET15b-*bldD*-Full, Amp$^{R}$ | (5) |
| pUZ8002 | RP4 derivative with defective oriT; Kan$^{R}$ | (9) |
| p3xFLAG | pIJ10170 derivative containing *3xFLAG* sequence downstream of MCS; Hyg$^{R}$ | (6) |
| pET15b | T7 expression vector; Amp$^{R}$ | Novagen |

| | | |
|---|---|---|
| pMAL-c2 | TAC expression vector, $Amp^R$ | NEB |
| pECJH1 | pET15b-*rmdB*; $Amp^R$ | This study |
| pECJH2 | pMAL-c2-*rmdA*; $Amp^R$ | This study |
| pECJH3 | pMAL-c2-*rmdA*$^{GGAAF}$; $Amp^R$ | This study |
| pECJH4 | pET15b-*rmdB*$^{GGDEF}$; $Amp^R$ | This study |
| pECJH5 | pET15b-*rmdB*$^{EAL}$; $Amp^R$ | This study |
| pSVNT28 | pET15b-*cdgE*; $Amp^R$ | This study |
| pSVJH02 | p3xFLAG *rmdB*-FLAG, $Hyg^R$ | (6) |
| pSVJH03 | p3xFLAG-*rmdB*$^{AAL}$, $Hyg^R$ | This study |
| pQE60 | pQE60-*pdeH*, C-terminal $His_6$ tag, $Amp^R$ | (3) |
| pRP89 | pET11- *pleD**, C-terminal His6 tag, $Amp^R$ | (4) |
| pIJ10663 | pET15b-*bldD*-Full | (5) |
| pSVSN-38 | pMS82-$_{ermE*}$-*chpC*, $Hyg^R$ | This study |
| pSVSN-39 | pMS82-$_{ermE*}$-*chpE*, $Hyg^R$ | This study |
| pSVSN-40 | pMS82-$_{ermE*}$-*chpH*, $Hyg^R$ | This study |
| pSVSN-65 | pMS82-$_{ermE*}$-*chpB*, $Hyg^R$ | This study |
| pSVSN-66 | pMS82-$_{ermE*}$-*chpD*, $Hyg^R$ | This study |
| pSVSN-67 | pMS82-$_{ermE*}$-*chpF*, $Hyg^R$ | This study |
| pSS5 | pIJ10750-$P_{ftsZ}$-*ftsZ-ypet*, $Hyg^R$ | (13) |
| SV-2_B03 | Cosmid, including *rmdA;* $Kan^R$, $Amp^R$ | JIC, Norwich, UK |
| SV-3_B05 | Cosmid, including *rmdB;* $Kan^R$, $Amp^R$ | JIC, Norwich, UK |
| SV-2_B03-$apr^R$_oriT | *kan::apr-oriT*; $Apr^R$, $Amp^R$ | This study |
| SV-3_B05-$apr^R$_oriT | *kan::apr-oriT*; $Apr^R$, $Amp^R$ | This study |

| | | |
|---|---|---|
| SV-2_B03-apr$^R$_oriT_ *rmdA*$^{ALLEF}$ | *kan::apr-oriT; rmdA::rmdA*$^{\mathrm{ALLEF}}$; Apr$^R$, Amp$^R$ | This study |
| SV-2_B03-apr$^R$_oriT_ *rmdA*$^{AAA}$ | *kan::apr-oriT; rmdA::rmdA*$^{\mathrm{AAA}}$; Apr$^R$, Amp$^R$ | This study |
| SV-3_B05-apr$^R$_oriT_*rmdB*$^{AAA}$ | *kan::apr-oriT; rmdB::rmdB*$^{\mathrm{AAA}}$; Apr$^R$, Amp$^R$ | This study |

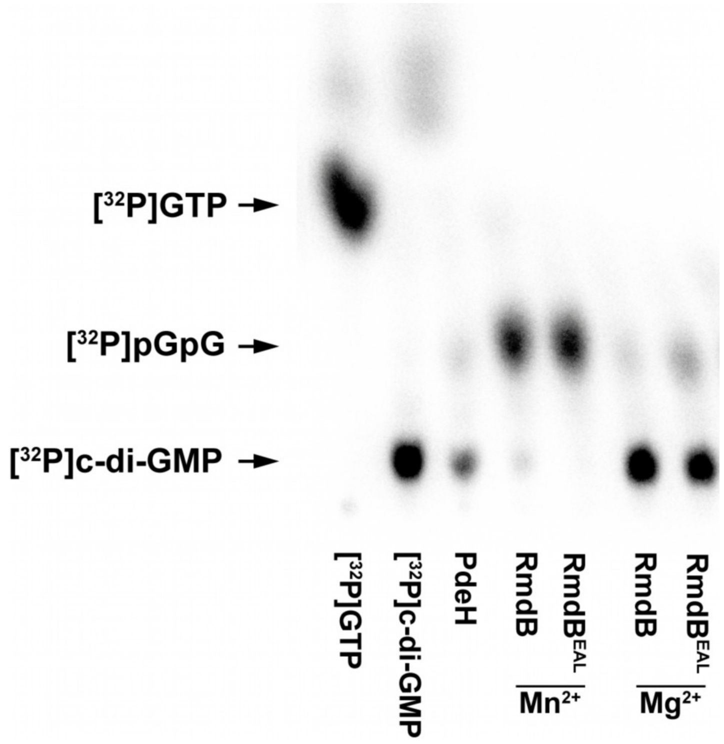


**Figure S1. RmdB and RmdB-EAL are more active in presence of $MnCl_2$ than $MgCl_2$.** The cytosolic fraction of RmdB and its isolated EAL domain (*rmdB*$_{421\text{-}704}$; RmdB$^{EAL}$) were expressed as N-terminally 6×His-tagged variants from pET15b in *E. coli* BL21 pLysS and purified using Ni-NTA based affinity chromatography. For growth of the overexpression strains, LB was supplemented with ampicillin (100 µg/ml) and either 0.35 mM $MgCl_2$ or $MnCl_2$. Proteins were eluted with elution buffer (50 mM Tris HCl, pH 8; 300 mM NaCl; 10% glycerol; 250 mM imidazole; 1 mM ß-mercaptoethanol) containing either 5 mM $MgCl_2$ or $MnCl_2$ and dialyzed twice against reaction buffer (25 mM Tris HCl, pH 7.5; 250 mM NaCl; 5 mM β-mercaptoethanol, 10% glycerol) with either 5 mM $MgCl_2$ or $MnCl_2$. One µM of RmdB and RmdB$^{EAL}$, respectively, was added to 2.08 nM [$^{32}$P]c-di-GMP as substrate and the reactions were separated by thin layer chromatography, exposed on a phosphor imaging screen and scanned by a phosphor imager.

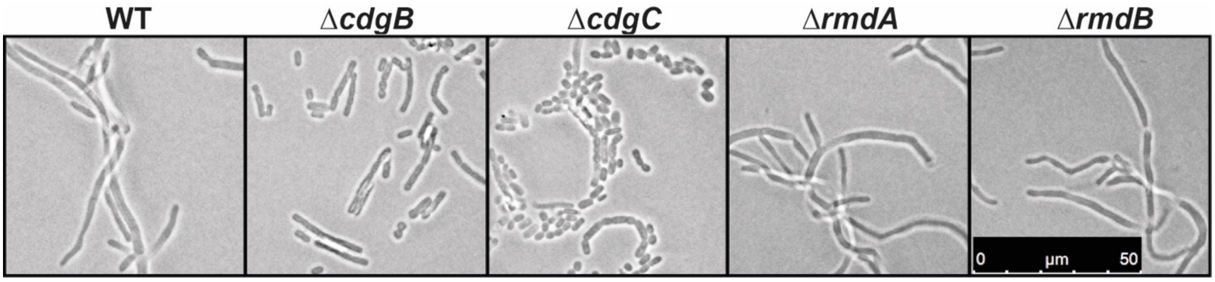


**Figure S2. Morphology of *S. venezuelae* wild type (WT), Δ*cdgB*, Δ*cdgC*, Δ*rmdA* and Δ*rmdB* strains when harvested for RNA-seq analysis.** Wild type, Δ*rmdA* and Δ*rmdB* grew vegetatively; Δ*cdgB* and Δ*cdgC* have already produced spores. Phase contrast microscopy images showing coverslip imprints from the upper layer of the macrocolonies grown for 30 h at 30 °C. Cells were imaged using the Leica DM2000 LED microscope at 100× magnification.

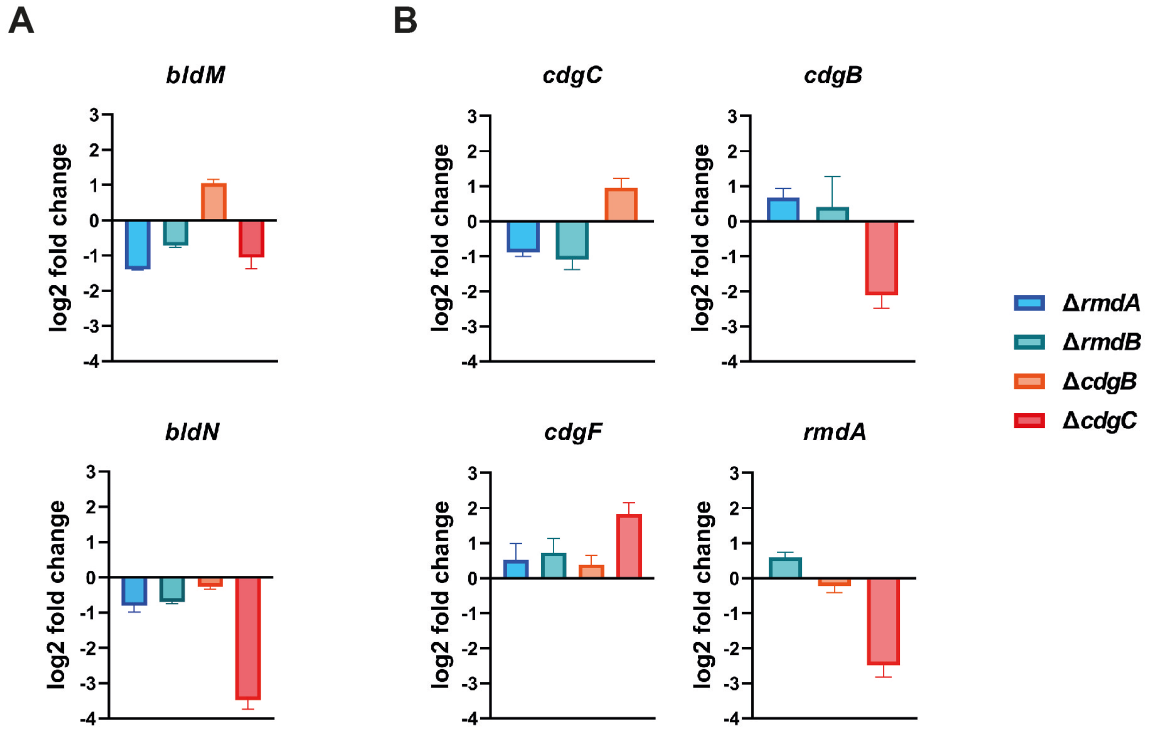


**Figure S3. qRT-PCR analysis of *bldM*, *bldN*, *cdgC*, *cdgB*, *cdgF* and *rmdA* expression in Δ*rmdA*, Δ*rmdB*, Δ*cdgB* and Δ*cdgC*.**

Strains were grown on Maltose-Yeast Extract-Malt Extract (MYM) plates for 30 h at 30 °C when harvested for RNA isolation using SV Total RNA Isolation Kit (Promega). qRT-PCR was performed using the SensiFAST SYBR No-ROX One-Step Kit (Bioline) and the CFX Connect real time system (Bio-Rad). The assay was reproduced at least three times with three technical replicates per experiment. Expression values were calculated relative to the accumulation of the constitutively expressed *hrdB* reference mRNA and normalized to the wild type value. Data are presented as mean of technical replicates ± standard deviation (n=3). log2 change >1/<-1 was considered as significant.

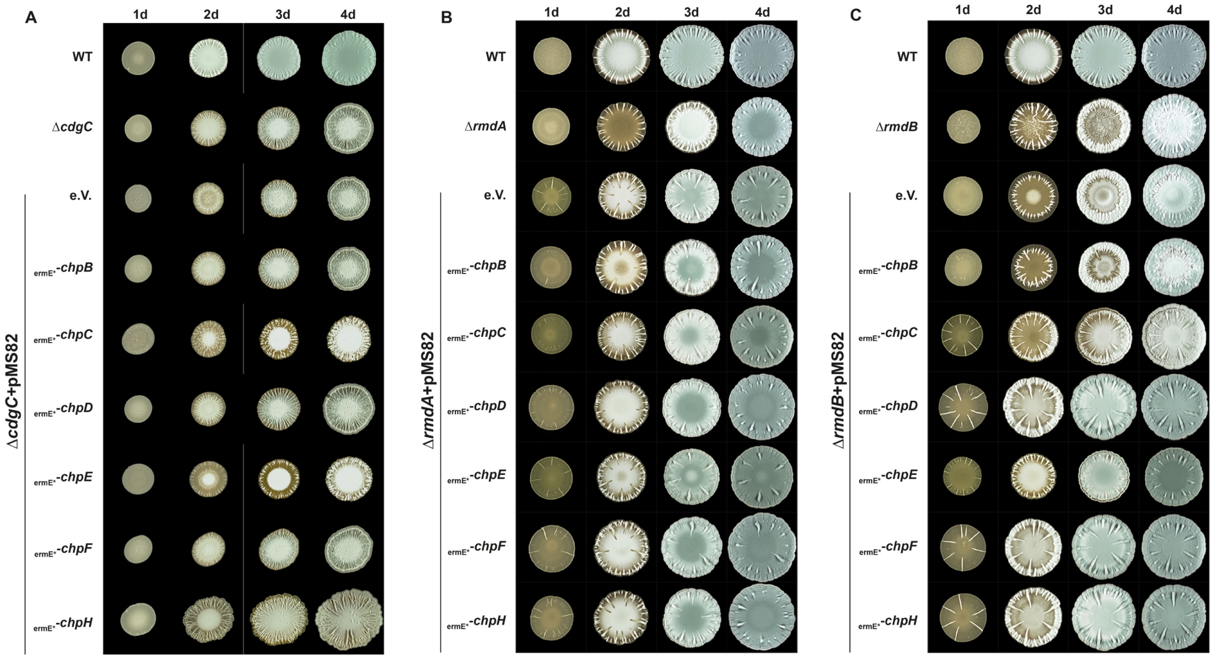


**Figure S4. Overexpression of chaplin genes in Δ*cdgC*, Δ*rmdA* and Δ*rmdB*.**

*chpB*-*F* and *chpH* were individually expressed from the integrative pMS82 vector controlled by the constitutive *ermE** promoter in Δ*cdgC* (A), Δ*rmdA* (B) and Δ*rmdB* (C). Twelve µl of $2x10^5$ CFU/µl *S. venezuelae* spores were pipetted on MYM agar and incubated for up to 4 days at 30 °C. Macrocolonies were photographed using a binocular (Stemi 2000C, Zeiss) equipped with a camera (AxioCAM ICc 3, Zeiss). e.V.: empty vector control.

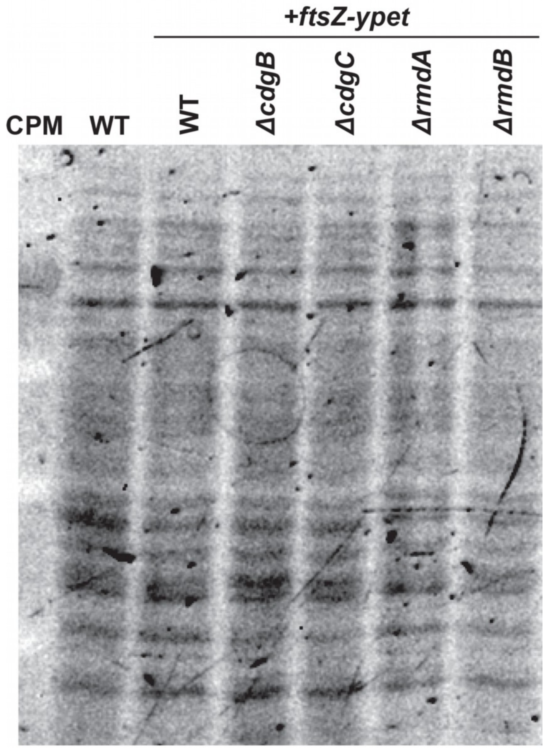


**Figure S5. Loading control for immunoblot analysis of FtsZ-YPet.**

*ftsZ-ypet* expressed from the integrative vector pSS5 in wild type, Δ*cdgB*, Δ*cdgC*, Δ*rmdA* and Δ*rmdB* strains that were grown in liquid MYM at 30°C for 12 h. Two ml samples were harvested, washed and resuspended as described in Schlimpert *et al.* 2017 (13). After resuspension in lysis buffer cells were homogenized using a BeadBeater (Biozym; six cycles at 6.00 m/s; 30 s pulse; 60 s interval) and cell debris was then centrifuged at 14.800 rpm for 10 min at 4°C. Total protein concentration was determined using Bradford assay (Roth) and adjusted to 1 mg/ml in each sample. Fourteen µg total protein per sample were loaded and separated on a 12 % SDS polyacrylamide gel via electrophoresis. For loading control, SDS polyacrylamide separation gels contained 0.5% 2,2,2-Trichloroethanol (TCE) that was added prior polymerization. After electrophoresis, gels were placed on a UV crosslinker (Stratagene) and the protein bands were treated with UV-light for 5 min to allow fluorescence detection on a 300-nm transilluminator.

### SUPPLEMENTARY REFERENCES


1. Huynh TN, Luo S, Pensinger D, Sauer JD, Tong L, Woodward JJ. 2015. An HD-domain phosphodiesterase mediates cooperative hydrolysis of c-di-AMP to affect bacterial growth and virulence. Proc Natl Acad Sci U S A 112:E747-56.
2. Christen M, Christen B, Folcher M, Schauerte A, Jenal U. 2005. Identification and characterization of a cyclic di-GMP-specific phosphodiesterase and its allosteric control by GTP. J Biol Chem 280:30829-37.
3. Pesavento C, Becker G, Sommerfeldt N, Possling A, Tschowri N, Mehlis A, Hengge R. 2008. Inverse regulatory coordination of motility and curli-mediated adhesion in *Escherichia coli*. Genes Dev 22:2434-46.
4. Paul R, Weiser S, Amiot NC, Chan C, Schirmer T, Giese B, Jenal U. 2004. Cell cycle-dependent dynamic localization of a bacterial response regulator with a novel di-guanylate cyclase output domain. Genes Dev 18:715-27.
5. Tschowri N, Schumacher MA, Schlimpert S, Chinnam NB, Findlay KC, Brennan RG, Buttner MJ. 2014. Tetrameric c-di-GMP mediates effective transcription factor dimerization to control *Streptomyces* development. Cell 158:1136-47.
6. Al-Bassam MM, Haist J, Neumann SA, Lindenberg S, Tschowri N. 2018. Expression Patterns, Genomic Conservation and Input Into Developmental Regulation of the GGDEF/EAL/HD-GYP Domain Proteins in *Streptomyces*. Front Microbiol 9:2524.
7. Hayashi K, Morooka N, Yamamoto Y, Fujita K, Isono K, Choi S, Ohtsubo E, Baba T, Wanner BL, Mori H, Horiuchi T. 2006. Highly accurate genome sequences of *Escherichia coli* K-12 strains MG1655 and W3110. Mol Syst Biol 2:2006 0007.
8. Herbst S, Lorkowski M, Sarenko O, Nguyen TKL, Jaenicke T, Hengge R. 2018. Transmembrane redox control and proteolysis of PdeC, a novel type of c-di-GMP phosphodiesterase. EMBO J 37.
9. Paget MS, Chamberlin L, Atrih A, Foster SJ, Buttner MJ. 1999. Evidence that the extracytoplasmic function sigma factor sigmaE is required for normal cell wall structure in *Streptomyces coelicolor* A3(2). J Bacteriol 181:204-11.
10. Datsenko KA, Wanner BL. 2000. One-step inactivation of chromosomal genes in *Escherichia coli* K-12 using PCR products. Proc Natl Acad Sci U S A 97:6640-5.
11. Thomason LC, Sawitzke JA, Li X, Costantino N, Court DL. 2014. Recombineering: genetic engineering in bacteria using homologous recombination. Curr Protoc Mol Biol 106:1 16 1-39.
12. Gust B, Challis GL, Fowler K, Kieser T, Chater KF. 2003. PCR-targeted *Streptomyces* gene replacement identifies a protein domain needed for biosynthesis of the sesquiterpene soil odor geosmin. Proc Natl Acad Sci U S A 100:1541-6.
13. Schlimpert S, Wasserstrom S, Chandra G, Bibb MJ, Findlay KC, Flärdh K, Buttner MJ. 2017. Two dynamin-like proteins stabilize FtsZ rings during *Streptomyces* sporulation. Proc Natl Acad Sci U S A 114:E6176-E6183.
